# Opposing macroevolutionary and trait-mediated patterns of threat and naturalization in flowering plants

**DOI:** 10.1101/2020.07.24.219667

**Authors:** J. P. Schmidt, T. J. Davies, M. J. Farrell

## Abstract

Due to expanding global trade and movement, new plant species are establishing in exotic ranges at increasing rates while the number of native species facing extinction from multiple threats grows. Yet, how species losses and gains globally may together be linked to traits and macroevolutionary processes is poorly understood. Here we show that, adjusting for diversification rate and age, the proportion of threatened species across flowering plant families is negatively related to the proportion of naturalized species. Moreover, naturalization is positively associated with climate variability, short generation time, autonomous seed production, and interspecific hybridization, but negatively with age and diversification; whereas threat is negatively associated with climate variability and hybridization, and positively with biotic pollination, age and diversification. Such a pronounced signature of naturalization and threat across plant families suggests that both trait syndromes have coexisted over deep evolutionary time and that neither strategy is necessarily superior to the other.

## Introduction

Ongoing habitat loss, degradation, and fragmentation (Corlett 2016), disruption of historic disturbance regimes (Alstad *et al*. 2016), increased invasion success of alien species (Winter *et al*. 2009), and climate change (Willis *et al*. 2008) are driving plant extinctions at accelerating rates (Alstad *et al*. 2016, Yessoufou & Davies 2016). At the same time, global trade has increased the pace at which alien plants are introduced and become established outside their native ranges (Seebens *et al*. 2017). Yet, while a broad body of work suggests threatened and invasive species contrast sharply in traits and have distinct phylogenetic distributions (Davies *et al*. 2011), our understanding of global patterns of threat and naturalization as potentially interrelated macroecological and macroevolutionary phenomena remains lacking (Vellend *et al*. 2017).

Extinction risk at broad scales appears to be strongly structured on plant phylogeny (*e.g*. Vamosi &Wilson 2008, Davies *et al*. 2011, Vamosi *et al*. 2018) with the frequency of threatened angiosperms highest within speciose clades (Schwartz & Simberloff 2001) and generally young, rapidly diversifying lineages (Davies *et al*. 2011). Critically, extinction risk may be more related to evolutionary dynamics (Davies *et al*. 2011) than to traits, and contingent on extinction drivers such as habitat loss, exploitation, etc., (Godefroid *et al*. 2014, Davies 2019). That extinction risk correlates strongly with clade age and richness suggests that the defining characteristics of rarity – endemism, limited range, and small population sizes (Rabinowitz *et al*.1986) – may simply follow from high rates of speciation (Davies *et al*. 2011). However, Vamosi and Wilson (2008) found extinction risk to be elevated in species-poor families, moreover, in some habitats, phylogenetically distinct species may be more threatened (Daru *et al*. 2013). Therefore, in old and species-poor families, perhaps remnants of formerly large and diverse clades, extinction risk may be linked to life history features that are sensitive to extinction drivers (Yessoufou & Davies 2016), while diversification dynamics dominate in young, species-rich families, implying that the drivers of extinction differ between old vs young clades (Vamosi *et al*. 2018, Davies 2019).

Successful plant invasions are conditioned on context-specific factors that include use (van Kleunen *et al*. 2020) and transport by humans (Kueffer 2017), degree of climate matching (Thuiller *et al*. 2005), residence time (Wilson *et al*. 2007), propagule pressure (Simberloff 2009), geography of habitat alteration and anthropogenic disturbance (Lembrechts *et al*. 2016), and the invasibility of particular communities and biogeographic regions (Richardson & Pyšek 2006). Nonetheless, successful invasions have been correlated in comparative studies with a suite of traits – auto-fertility (Razanajatovo *et al*. 2016), self-compatibility (Hao *et al*. 2011), height (van Kleunen *et al*. 2007), small seeds (Hamilton *et al*. 2005), high specific leaf area (Hamilton *et al*. 2005), large native range size (Schmidt *et al*. 2012, van Kleunen *et al*. 2007), broad climate and habitat tolerances (Schmidt *et al*. 2012, Bradshaw *et al*. 2008), competitive ability (Guo *et al*. 2018), variability and perhaps plasticity in growth form and life history (Schmidt *et al*. 2012), abiotic pollination (Hao *et al*. 2011), polyploidy (Schmidt *et al*. 2012) and hybridization (Ellstrand & Schierenbeck 2000) – that appear consistently advantageous (Table 1). Thus, while naturalized species may also be non-randomly distributed among angiosperm families (Pyšek 2017, Pyšek 1998) and benefit from factors extrinsic to ecological features, trait syndromes across clades appear to strongly influence invasion success.

Here, adjusting for phylogenetic relatedness and clade size, we ask 1) how the representation of globally threatened species by flowering plant family is related to that of naturalized species (Fig. 1); 2) whether the frequency of species in either category tends toward the opposite extremes of the same trait axes; and 3) if the relative importance of traits vs evolutionary history as explanatory variables differs for threat vs naturalization. We evaluate these hypotheses in a hierarchical Bayesian regression framework – allowing us to quantify uncertainty in analyses, and to better compare effect sizes among predictors.

**Fig. 1.**
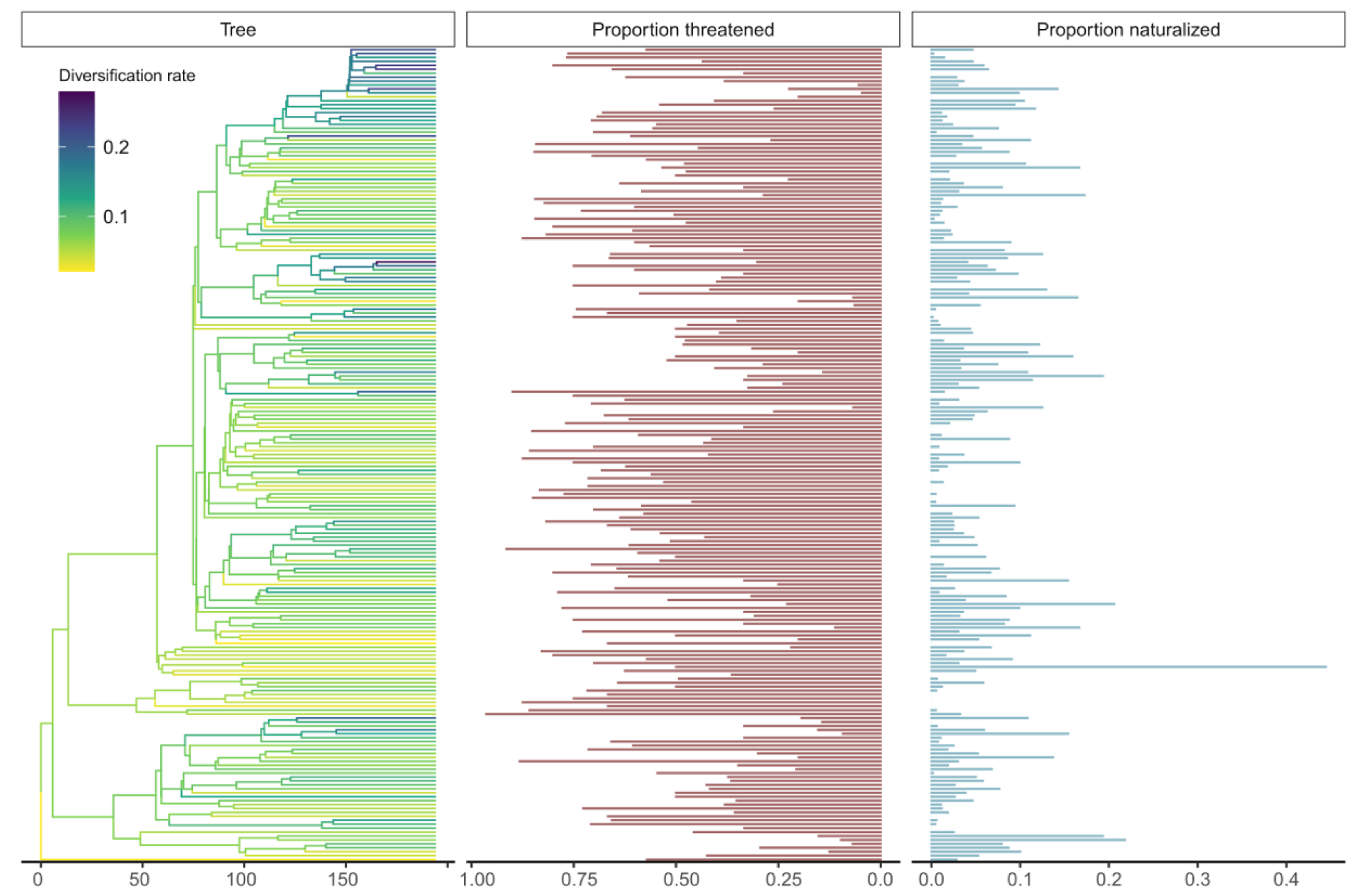
Phylogeny of the 207 angiosperm families. Families included have at least one species vetted by the IUCN and less than 100% of species categorized as either threatened or naturalized. Internal branches (*left*) show family level diversification rate, log(species richness)/clade age (ancestral state reconstruction calculated using *fastAnc* in the phytools R package). Proportion IUCN vetted species classed as globally threatened in red (*middle*), and proportion of all species in the family naturalized in blue (*right*).

## Results

Across angiosperm families, the proportion of vetted species classed as threatened was negatively related to the frequency of naturalized species (Fig. 2). In separate models predicting the proportion by family of threatened to vetted species or naturalized to total species as a function of traits and evolutionary history, the proportion threatened was positively related and the proportion naturalized negatively related – to family age and diversification rate (Fig. 3). Trait covariates also showed opposing patterns for threatened and naturalized models. Threat was lower in families with greater climate variability (number of climate zones occupied), and those characterized by many interspecific hybrids and predominantly herbaceous life forms, and higher in families that included animal pollinated species (Fig. 3). In contrast, proportion naturalized per family was positively related to herbaceous growth form, climate variability, number of interspecific hybrids, and the presence of species with asexual seed production, and annual life-histories, but negatively correlated with animal pollination (Fig. 3). Notably, the negative effect sizes of diversification rate and family age on proportion naturalized were greater in magnitude than the positive effects of these same factors on proportion threatened (Fig. 3).

**Fig. 2.**
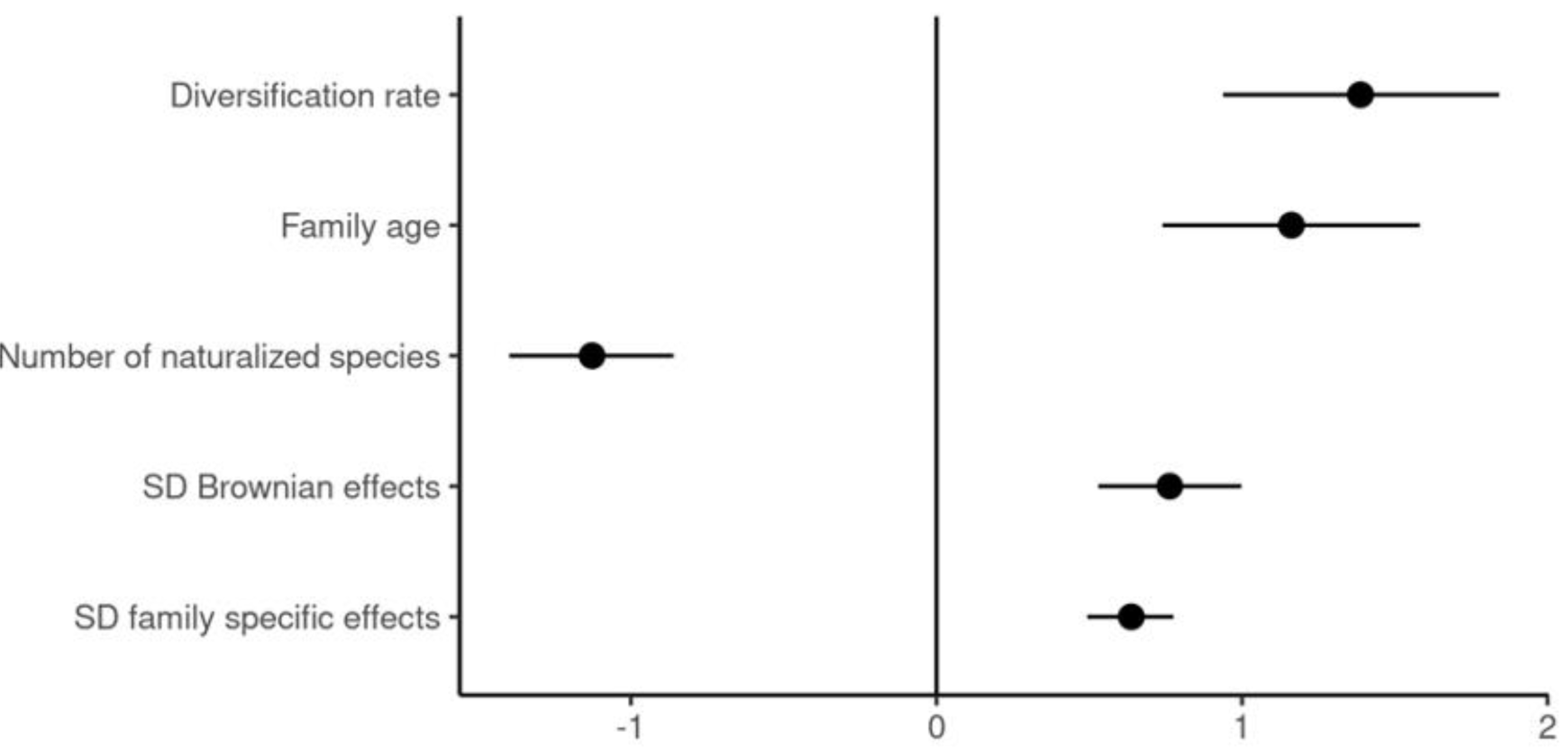
Estimated model parameters for predicting proportion threatened. Regression coefficients and 80% credible intervals for hierarchical effects by family among the 236 families with IUCN vetted species as a function of the number of naturalized species, while adjusting for diversification rate (log(family size)/family age) and family-level effects.

**Fig. 3.**
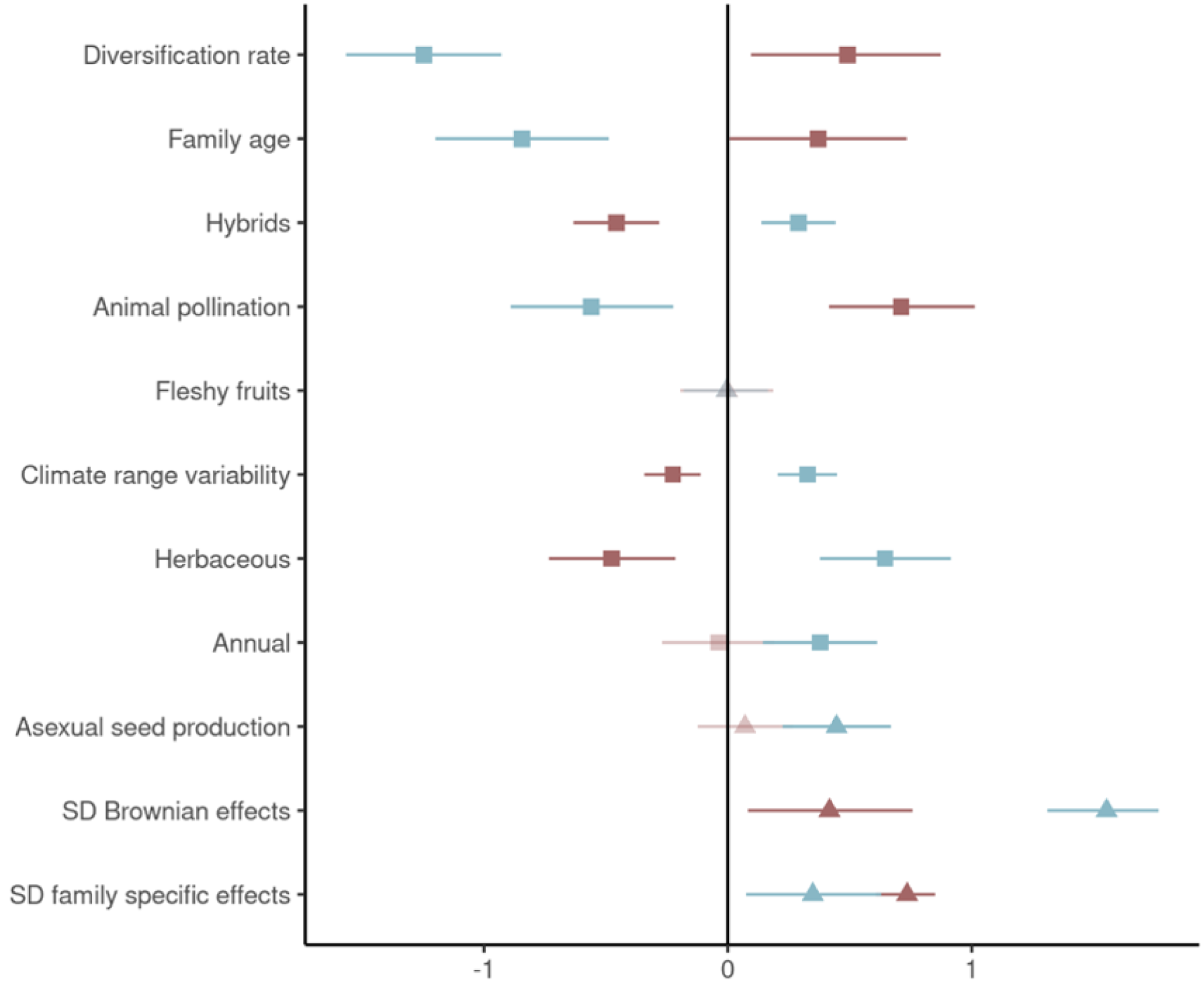
Estimated model parameters for threatened and invasive. Regression coefficients and hierarchical standard deviations, with 80% credible intervals, by family (*red*) among the 267 families with IUCN vetted species and the proportion naturalized (*blue*) among all species per family for all 395 families included in the study. Squares (rather than triangles) indicate variables with opposite effects on naturalized vs threatened status, pale triangles credible intervals overlapping zero.

Model fit (normalized root mean squared error: *NRMSE*) for threat (*NRMSE* = 0.20, *sd* = 0.03) was better than for naturalization (*NRMSE* = 0.54, *sd* = 0.08). And, contrary to expectations, the proportion of the variance in family level effects explained by the Brownian phylogenetic component (phylogenetic heritability, Lynch’s *λ*), was higher for naturalization (*λ* = 0.93, s*d* = 0.08) than threat (*λ* = 0.25, *sd* = 0.22, Fig. 4). Sensitivity analyses using the total number of species per family rather than IUCN vetted species in the binomial response (*NRMSE* = 0.49, *sd* = 0.06, Tables S3,S4) and excluding family level effects (*NRMSE* = 0.55, *sd* = 0.03, S3.1) both reduced threat model fit, demonstrating the importance of controlling for data deficiencies and family level variation not accounted for by trait data. Sensitivity analyses investigating the proportion of invasive rather than naturalized species per family also reduced model fit (*NRMSE* = 0.78, *sd* = 0.11) likely reflecting the smaller number of taxa classed as invasive, and perhaps the more arbitrary application of the invasive vs naturalized label.

**Fig. 4.**
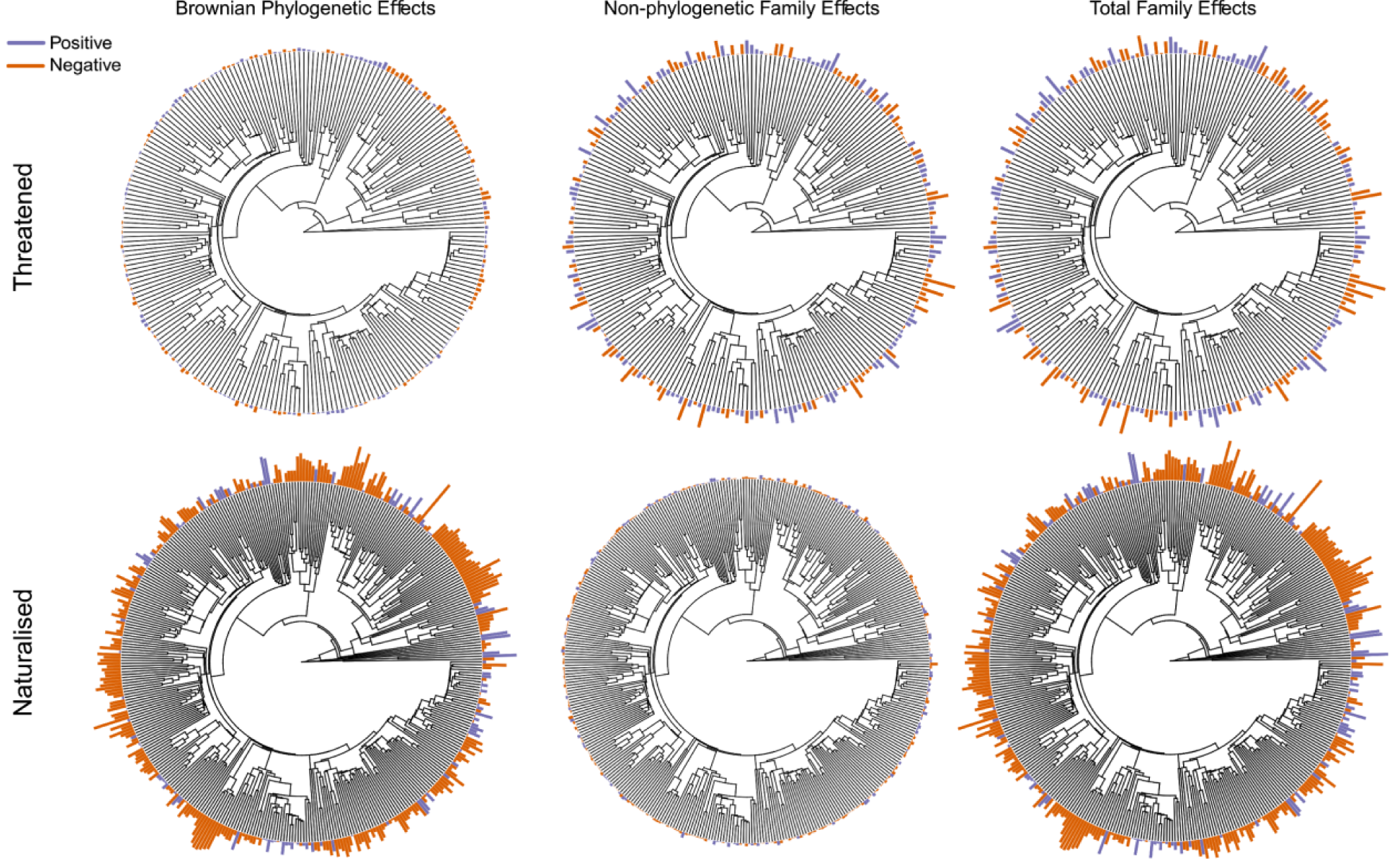
Family-level hierarchical effects for the full model. Proportion IUCN vetted species threatened per family (*top row*), and the proportion naturalized (*bottom row*) among all species per family, separated into Brownian phylogenetic, non-phylogenetic, and total family effects (Brownian + non-phylogenetic). Purple bars indicate positive effects, orange bars indicate negative effects, and bar lengths indicate the relative magnitude of the mean estimated effect per family.

## Discussion

At the global scale, we show that the drivers of threat and naturalization across angiosperms are opposing, and that this can be explained by divergent macro-evolutionary and ecological trait relationships. We note that models excluding family-specific effects or phylogeny were poorer fits, indicating that a large component of the variation among families still remains unexplained; and family-level estimates of age and diversification rate (not entirely separable) may not optimally capture the signature of macroevolutionary processes towards the present. Nonetheless, our results provide the first quantitative support at a global scale across angiosperms for the hypothesis that naturalization and threat represent “two sides of the same coin” (Schmidt *et al*. 2012, Bradshaw *et al*. 2008).

Consistent with previous studies (Davies *et al*. 2011), diversification rate was positively related to proportion threatened per family. In contrast, naturalization was negatively related to both diversification rate and family age. Fast diversifying clades are often associated with localized radiations that give rise to many endemics with narrow ranges that are restricted to particular habitat types (*e.g*. many plant lineages in the Fynbos, South Africa, have diversified rapidly, and are characterized by a high diversity of narrowly distributed species, frequently restricted to particular soil types). This high ecological specialization likely restricts geographical expansion and naturalization outside of the native range, resulting in a negative correlation between diversification and naturalization. More established lineages, and those in older families may have had more time to spread and thus have had more opportunities to become naturalized outside of their native range. However, as species age they may also contract in their geographic extent, especially if the environmental conditions which favored their initial expansion change (c.f. taxon cycle: Ricklefs & Bermingham 2002). Species experiencing range contraction are also less likely to become naturalized elsewhere. As only old families can contain old species, this could lead to a negative correlation between naturalization and family age. The magnitude of the negative effects of diversification rate and family age on naturalization were also somewhat greater than those of the positive effects on threat. Contrary to initial expectations, after adjusting for macroevolutionary and life-history predictors, we also found that in the naturalization model, a larger fraction of variance in family level effects was attributable to Brownian phylogenetic effects, compared to the equivalent model for threat. We suggest that one possible explanation for the weaker signature of macroevolutionary process on threat is the mixing of threatened taxa found within both young and old clades such that observed threat captures two independent processes.

The opposing relationship between threat and naturalization was also manifested in the trait syndromes that characterize either status. Naturalization was positively related to herbaceous growth form, climate variability, frequency of interspecific hybrids, asexual seed production, and annual life history, and negatively related to animal pollination; whereas threat was negatively related to herbaceous growth form, climate variability, and frequency of interspecific hybrids, and positively to animal pollination and variation in growth form. The correlations we detect among angiosperm families match closely to expectations from theory and existing comparative studies across various scales (Table 1) – with naturalization characterized by habitat/climatic generalism, short generation time, asexuality and polyploidy, and threat by habitat specialism and endemism, dependence on mutualists, and diploidy.

That we recover such strong associations at higher taxonomic levels and at a global scale is notable and suggests that the trait signatures characterizing threat and naturalization in the present day extend back over deep evolutionary time. Thus, while particular trait syndromes appear to predispose some species to higher risk of extinction and others to ecological expansion, this might not translate straightforwardly to long-term macroevolutionary dynamics. For example, the traits characteristic of threatened species today may be features that permit chronically rare species to persist (*e.g*. the directed movement of pollen by an animal vector) and, by limiting outcrossing, allow adaptation to spatially restricted environmental conditions (Vermeij & Grosberg 2018). An alternative strategy – promoting colonization and expansion at range margins – relies on features such as abiotic pollination, asexual seed production, and, to enable rapid niche shifts in the face of novel climatic conditions or sudden environmental change, hybridization and polyploidy (Baniaga *et al*. 2019). Both strategies thus represent successful macro-evolutionary syndromes, but under contrasting selective regimes. Include a Discussion that summarizes (but does not merely repeat) your conclusions and elaborates on their implications. There should be a paragraph outlining the limitations of your results and interpretation, as well as a discussion of the steps that need to be taken for the findings to be applied. Please avoid claims of priority.

## Materials and Methods

### Data

To estimate species richness, we tabulated the number of species (including interspecific hybrids) within each of 404 angiosperm families (per APG3, Angiosperm Phylogeny Group 2009) from The Plant List (TPL, http://www.theplantlist.org/) – counting only the 301,639 species with ‘accepted’ names. To determine the number of species in each class, we compiled a list of all angiosperms labeled as 1) naturalized (12,256 species) in any region, globally, from the GloNAF database (van Kleunen 2019); 2) ‘invasive’ (4,540 species) from Randall (2017); and 3) as ‘threatened’ (12,894 species), if listed as globally “Vulnerable” or more threatened by the International Union for the Conservation of Nature (https://www.iucnredlist.org/, accessed March 11, 2019). We labeled species as ‘not threatened’ if listed as ‘Least Concern’, ‘Near Threatened’ or ‘data deficient’ by the IUCN. To control for study effort, we summed the number of species per family that have currently been assessed by the IUCN. In all cases (naturalized, invasive, threatened and vetted), we filtered species through TPL to include only accepted names. Of the 404 families, 239 included a naturalized species, 196 included an invasive species, 237 included an IUCN-vetted species, and all vetted families included a threatened species.

For family level trait data, we retrieved binary data on climate zones (tropical, subtropical, temperate, frigid zone) occupied, and presence of animal pollination and fleshy fruits – from the Watson and Dallwitz (1992 onward) online key to angiosperm families. To capture variability in climate tolerance, we summed the number of climate zones (1-4) reported among species within each family. Animal pollination and fleshy fruits were coded for each family as 1 if biotic pollination (346 families) or fleshy fruits (77 families) was the sole mode, 0.5 if multiple traits (biotic and abiotic pollination (13 families) or fleshy and non-fleshy fruits, 130 families), and 0 if only abiotic pollination (45 families) or dry fruits (197 families) are known. From Watson and Dallwitz and data from the Plants National Database (https://plants.sc.egov.usda.gov/), we labeled families as known to include annual species (1, 99 families) or otherwise (0, 305 families). From Zanne *et al*. (2014) and Hawkins *et al*. (2011) we classified families as predominantly herbaceous (1, 156 families), predominantly woody (0, 192 families), or mixed (0.5, 56 families). Lastly, we used data from Hojsgaard *et al*. (2014) to identify families in which asexual seed production (1, 173 families) is known to occur vs all other families (0, 231 families).

For estimates of age and diversification rates, we relied on stem family ages from the Zanne *et al*. (2014) phylogeny of vascular plants. Zanne *et al*. first used a method of congruification to make their tree ultrametric based on the phylogeny of Soltis *et al*. (2011). Branch lengths were then time-scaled using 39 fossil calibration points, which at the time represented the most reliable set of fossils spanning the angiosperm phylogeny.

### Statistical Analysis

We used Bayesian binomial-logit multilevel regressions to model the proportion of species threatened and naturalized plants per family. To assess the relationship between threat and naturalization, we fit an initial model with, for each family, the proportion of all species naturalized, and family age and diversification rate as continuous predictors, and with the proportion of vetted species (those assessed by the IUCN) that have been classed as threatened as the response. To aid in the comparison of effect sizes across continuous and binary predictors, we log-transformed, centered and scaled to a standard deviation of 0.5 all continuous predictors prior to analyses. To identify traits associated with the proportion of species threatened and naturalized per family, we fit two additional models that included family-level traits, and family age and diversification rate. We calculated diversification rate as log(*N*)/clade age, which is the maximum likelihood estimate of diversification rate assuming negligible extinction (eq. 3 in Magallon & Sanderson 2001). A more complex formulation (eq. 6 in Magallon & Sanderson 2001) allows a constant extinction, but empirically, family level diversification rates estimates are little influenced (Jansson & Davies 2008). To account for phylogenetic non-independence and family-level effects in all models, we included hierarchical effects by family separated into phylogenetic (assuming a Brownian motion model of evolution) and non-phylogenetic effects, following the additive quantitative genetic model adapted to interspecific data. The correlation matrix for determining family-level Brownian phylogenetic effects was calculated from the Zanne *et al*. (2014) phylogeny of vascular plants pruned to a single representative species per family.

We fit all models in Stan version 2.18.0 (The Stan Development Team) accessed using the R package *brms* version 2.7.0 (Buerkner 2017). We fit models with the *brms* default priors, which use a uniform prior for the regression coefficients, and half-Student’s *t* distributions with three degrees of freedom and scale parameter of 10 for the variance components of the family-level effects. As these represent extremely weak priors, for the simplified and full threatened models we also explored the use of alternative custom priors (normal (0,1.5) for regression coefficients and normal (0,1) for family effect standard deviations), which provide more realistic expectations (Figs. S1, S2). We found a negligible influence of each set of priors on posterior estimates (Figs. S4, S5, S7, S8, S10, S11, S13, S14), and thus assumed the *brms* default priors for all other models. We ran models across four chains, with 10000 iterations per chain. For each chain, the first 5000 iterations were used as burn-in and discarded. The remaining iterations were thinned to retain every fourth iteration, resulting in a total of 5000 posterior draws. We diagnosed convergence by visual inspection of traceplots and observation of Rhat values equal to 1.0 for all estimated parameters, and we assessed model fit using posterior predictive checks (see Data & Code supplement) and root mean square error (*RMSE*). To compare model fits across varying responses, we calculated the normalized *RMSE* (*NRMSE*) scaling by the interquartile range of the observed data, and across all posterior samples. To calculate phylogenetic heritability (Lynch 1991) we used the *hypothesis* function in *brms* with the hypothesis σ_2Brownian_ /(σ_2brownian_ + σ_2family-specfic_) = 0 across all posterior samples.

To explore the effect of including both Brownian and family-specific effects, we ran sensitivity models which fit the full threatened model with a) no family specific effects, b) only family specific effects, b) only Brownian effects. To determine the effect of using the number of IUCN vetted species rather than the total number of species per family, we fit the full threatened model with total species richness per family as *n* (the denominator) in the binomial model. Finally, to assess the sensitivity of the naturalized model results to the definition of an invasive species, we re-ran the naturalized model using the number of invasive species, defined by Randall (2017) as naturalized species that have spread and have significant ecological/economic impact.

## Funding

We thank the Macroecology of Infectious Disease Research Coordination Network, funded by NSF (DEB 1316223), for facilitating discussion among the authors, and for supporting MJF as a postdoctoral research associate.

## Author contributions

JPS, MJF, and TJD designed the study. JPS compiled the data. MJF conducted the analyses. JPS wrote the manuscript with input from TJD and MJF. The authors declare no conflicts of interest.

## Competing interests

Include any financial interests of the authors that could be perceived as being a conflict of interest. Also include any awarded or filed patents pertaining to the results presented in the paper. If there are no competing interests, please state so.

## Data and materials availability

All data and code necessary to reproduce the results are included in https://figshare.com/s/6d032d41cf669f4bd6e1 (reserved DOI: 10.6084/m9.figshare.11372037).

Opposing macroevolutionary and trait-mediated patterns of threat and naturalization in flowering plants

## Supplementary Materials

### 1 Exploration of priors

As a default for binomial-logit models, the *brms* package uses extremely flat priors for the regression coefficients and standard deviation parameters. These are improper unbounded uniform priors, and a half Student t distribtion with three degrees of freedom and a scale parameter of ten, respectively. To explore the influence of these priors, and their impact on model performance, we compare them to custom priors reflecting recommendations by Richard McElreath in his textbook *Statistical Rethinking* (a normal(0,1.5) and half-normal(0,1) respectively).

**Figure SM 1:**
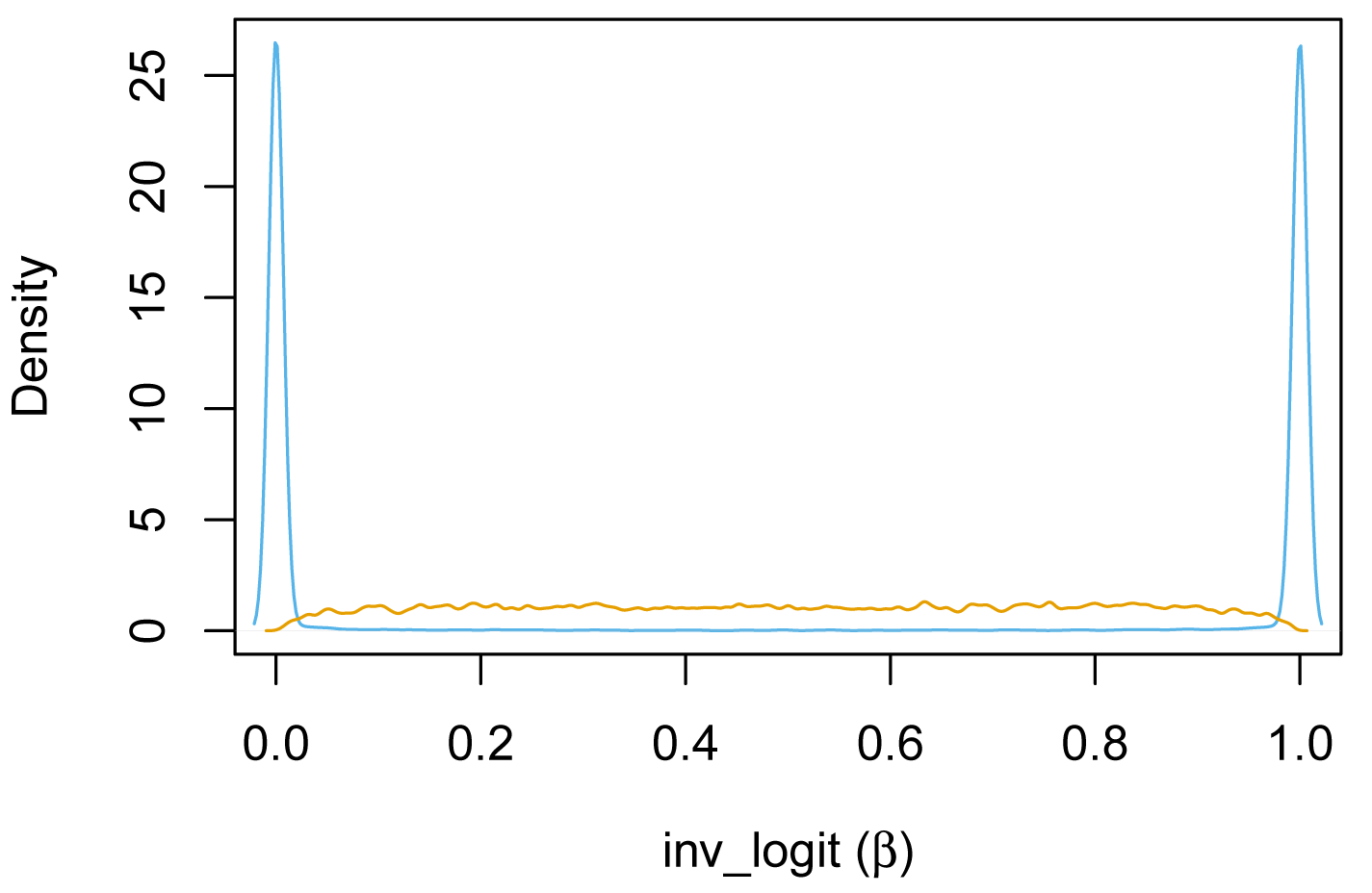
Priors on the regression coefficients for the *brms* default (blue), and custom (yellow) priors

The default improper uniform prior used by *brms* is flat on the log-odd scale, so when converted to the probability scale, puts excess mass near 0 and 1. A Normal(0, 1.5) prior is more reasonable, having a flatter distribution with a mode at 0.5 on the probability scale.

**Figure SM 2:**
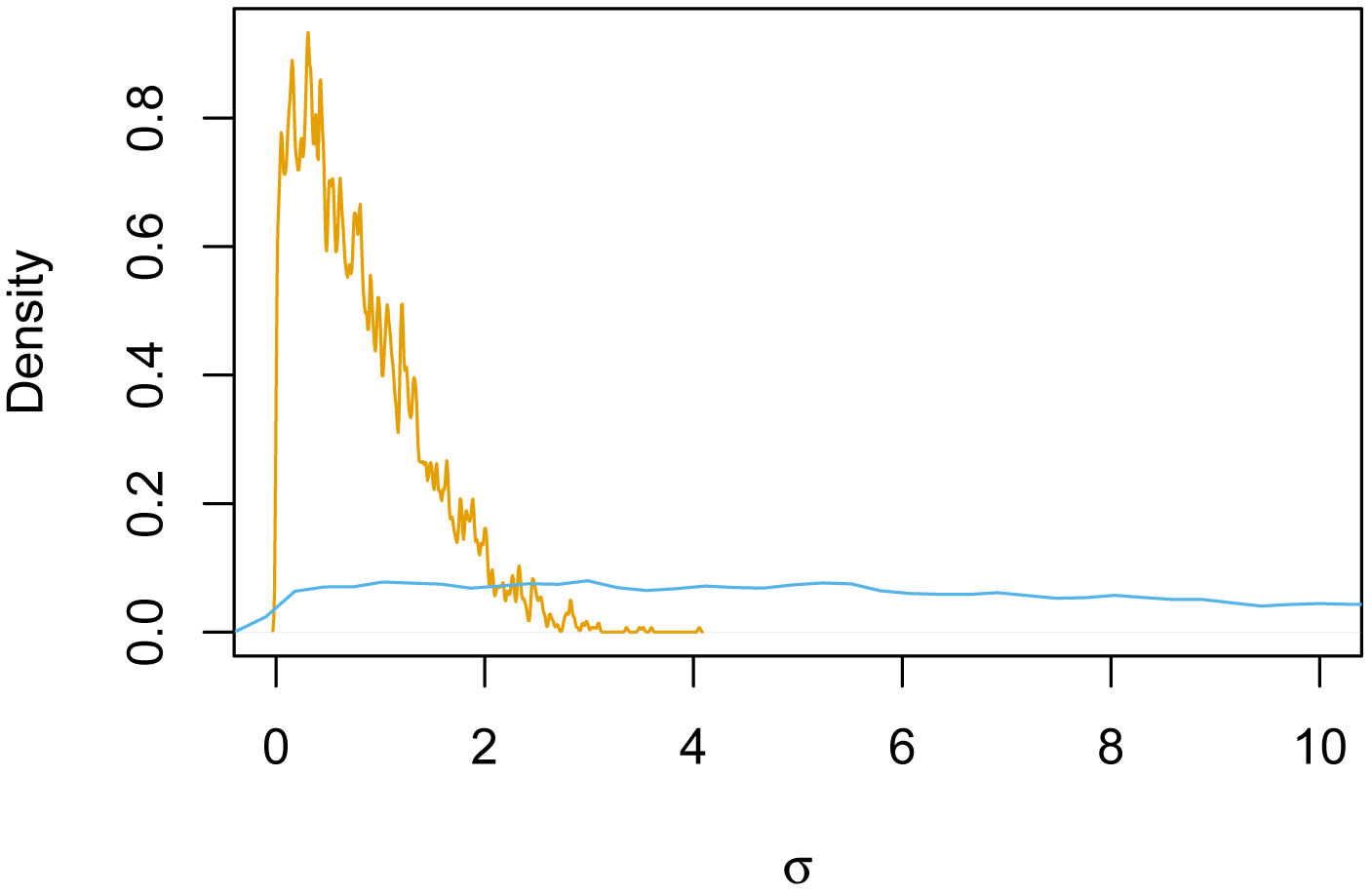
Priors on the regression coefficients for the *brms* default (blue), and custom (yellow) priors

The *brms* prior is extremely long tailed. Considering we are fitting a non-linear model in which changes in log-odds over over 4 have a diminishing influence due to the ceiling effect of the binomial distribution. With this in mind, a prior of Normal(0,1) constrains the posterior to a more realistic range, and is likely to improve sampling efficiency.

To explore the impact of each of these priors, we fit the simple and full threatened models using each set of priors, and visualize the relationships between the priors and associated posteriors for two representative parameters. Overall, the choice of prior was found to have no influence on the posterior parameter estimates in these two models, so we used the brms defaults for all other models.

### 2 Main models

#### 2.1 Simple threatened model - *brms* default priors

**Table SM 1:** Summary of model output for *β* and *σ* parameters, including posterior means, posterior standard deviations, 2.5%, 50%, and 97.5% quantiles, the effective sample size (n eff), and the potential scale reduction statistic (Rhat). n=236; RMSE=5.95 (+/-0.73 SD); NRMSE = 0.20 (+/-0.03).

|  | mean | sd | 2.5% | 50% | 97.5% | n_eff | Rhat |
| --- | --- | --- | --- | --- | --- | --- | --- |
| Number of naturalized species | -0.97 | 0.20 | -1.37 | -0.97 | -0.56 | 4320.48 | 1.00 |
| Diversification rate | 1.67 | 0.35 | 1.00 | 1.66 | 2.36 | 4526.64 | 1.00 |
| Family age | 1.33 | 0.33 | 0.68 | 1.33 | 1.98 | 4473.99 | 1.00 |
| SD brownian effects | 0.98 | 0.20 | 0.59 | 0.98 | 1.37 | 991.53 | 1.00 |
| SD family specific effects | 0.60 | 0.14 | 0.28 | 0.61 | 0.83 | 813.83 | 1.00 |

**Figure SM 3.**
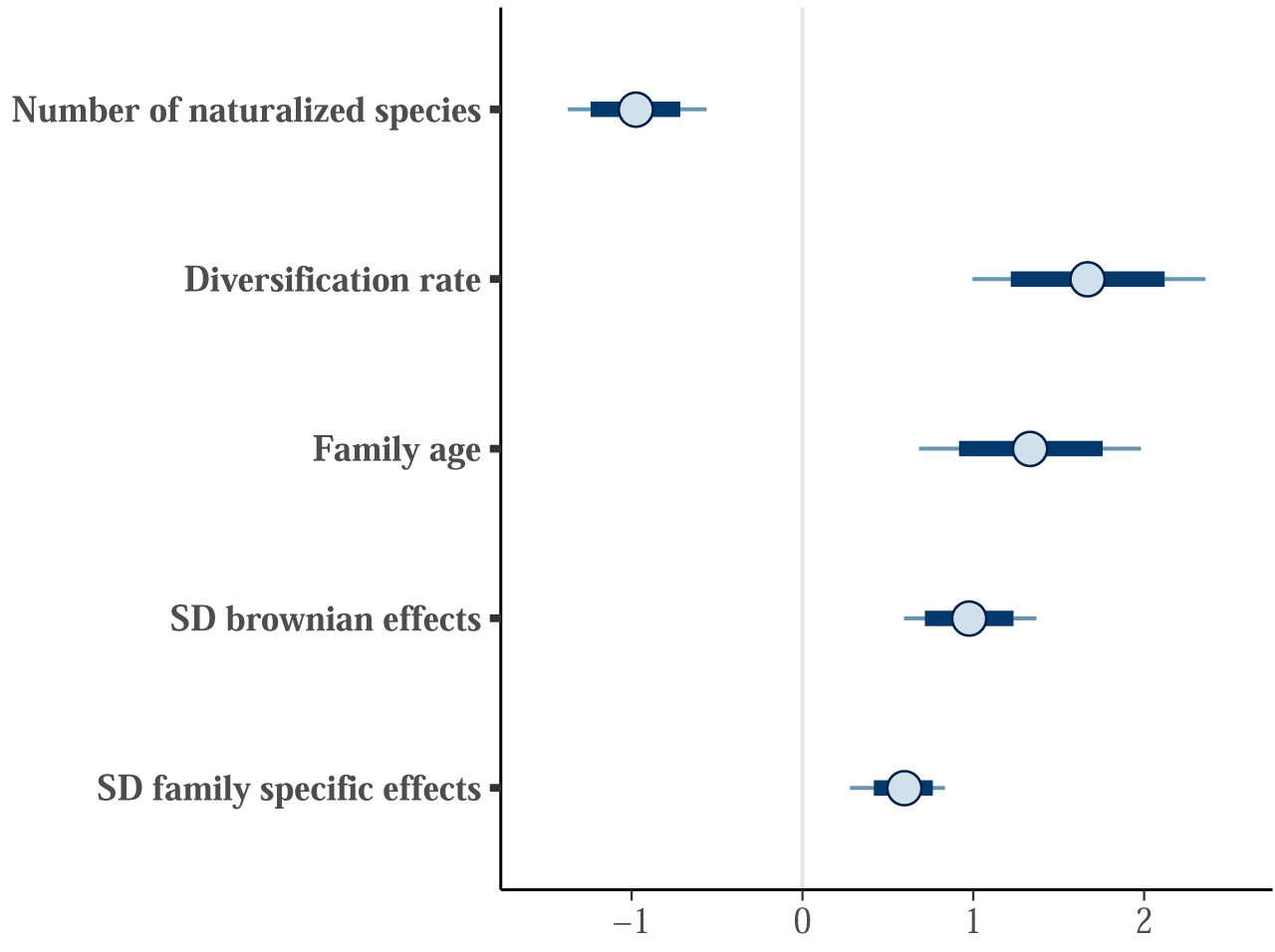
Forest plot for estimated *β* and *σ* parameters for the simple threatened model. Points represent posterior means, with thick lines represeting 80% credible intervals, and thin lines representing 95% credible intervals.

**Figure SM 4:**
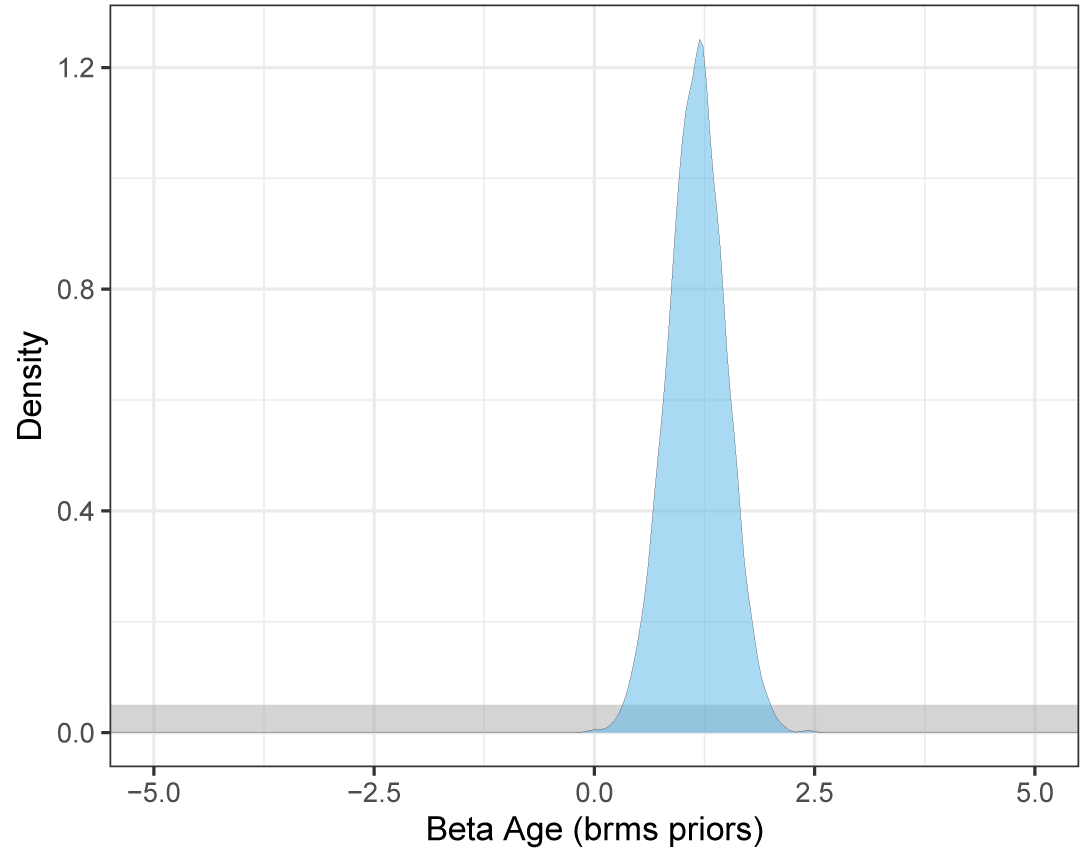
Prior (grey) and posterior (blue) distributions for the Age regression coefficient for the simple rarity model using *brms* priors. The improper uniform prior was restricted to (−10,10) to aid visualization.

**Figure SM 5:**
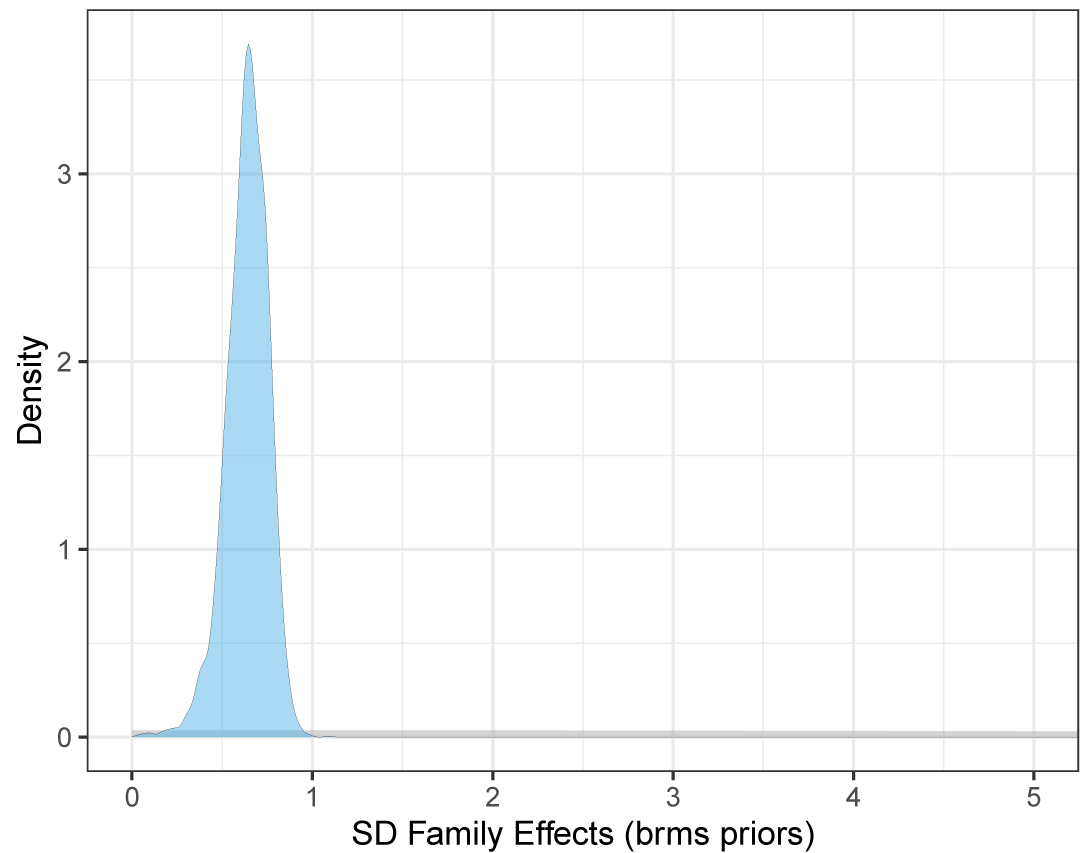
Prior (grey) and posterior (blue) distributions for the Family Effects standard deviation parameter for the simple rarity model using *brms* priors.

### 2.2 Simple threatened model - custom priors

**Table SM 1:** Summary of model output for *β* and *σ* parameters, including posterior means, posterior standard deviations, 2.5%, 50%, and 97.5% quantiles, the effective sample size (n eff), and the potential scale reduction statistic (Rhat). n=236; RMSE=5.93 (+/-0.73 SD); NRMSE = 0.20 (+/-0.03).

|  | mean | sd | 2.5% | 50% | 97.5% | n_eff | Rhat |
| --- | --- | --- | --- | --- | --- | --- | --- |
| Number of naturalized species | -0.91 | 0.20 | -1.29 | -0.91 | -0.52 | 4462.00 | 1.00 |
| Diversification rate | 1.52 | 0.33 | 0.87 | 1.52 | 2.16 | 4579.12 | 1.00 |
| Family age | 1.20 | 0.32 | 0.57 | 1.20 | 1.82 | 4506.43 | 1.00 |
| SD brownian effects | 0.95 | 0.19 | 0.59 | 0.95 | 1.33 | 1175.25 | 1.00 |
| SD family specific effects | 0.61 | 0.13 | 0.30 | 0.62 | 0.83 | 1044.40 | 1.00 |

**Figure SM 6:**
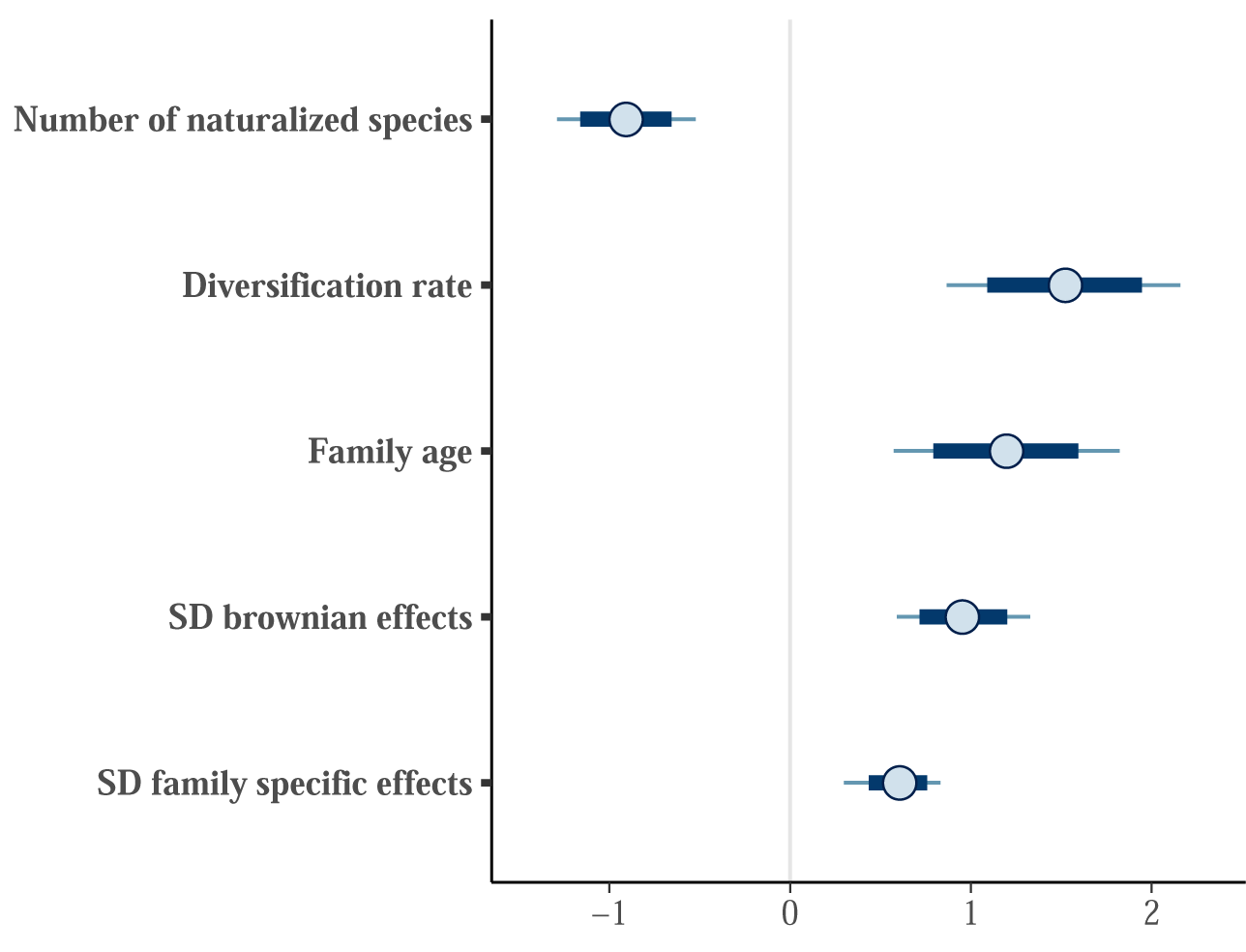
Forest plot for estimated *β* and *σ* parameters for the simple threatened model with custom priors. Points represent posterior means, with thick lines represeting 80% credible intervals, and thin lines representing 95% credible intervals.

**Figure SM 7:**
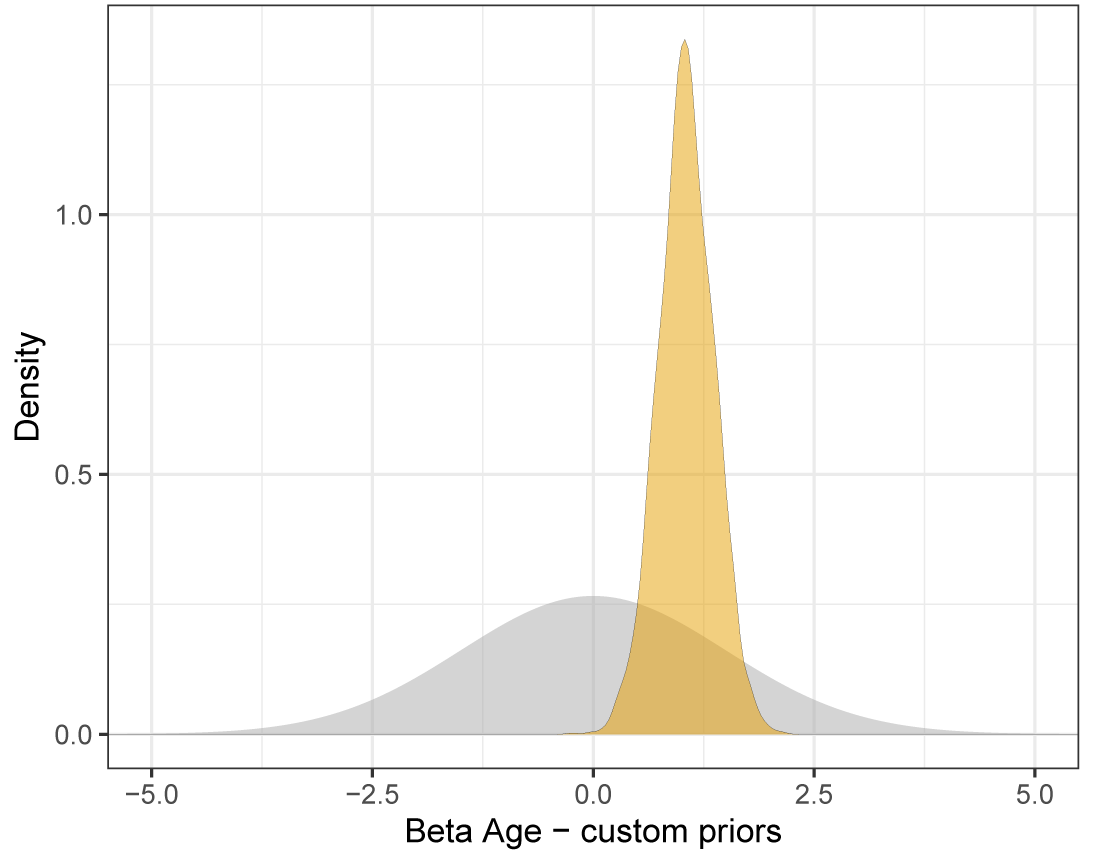
Prior (grey) and posterior (yellow) distributions for the Age regression coefficient for the simple rarity model using custom priors. The improper uniform prior was restricted to (−10,10) to aid visualization.

**Figure SM 8:**
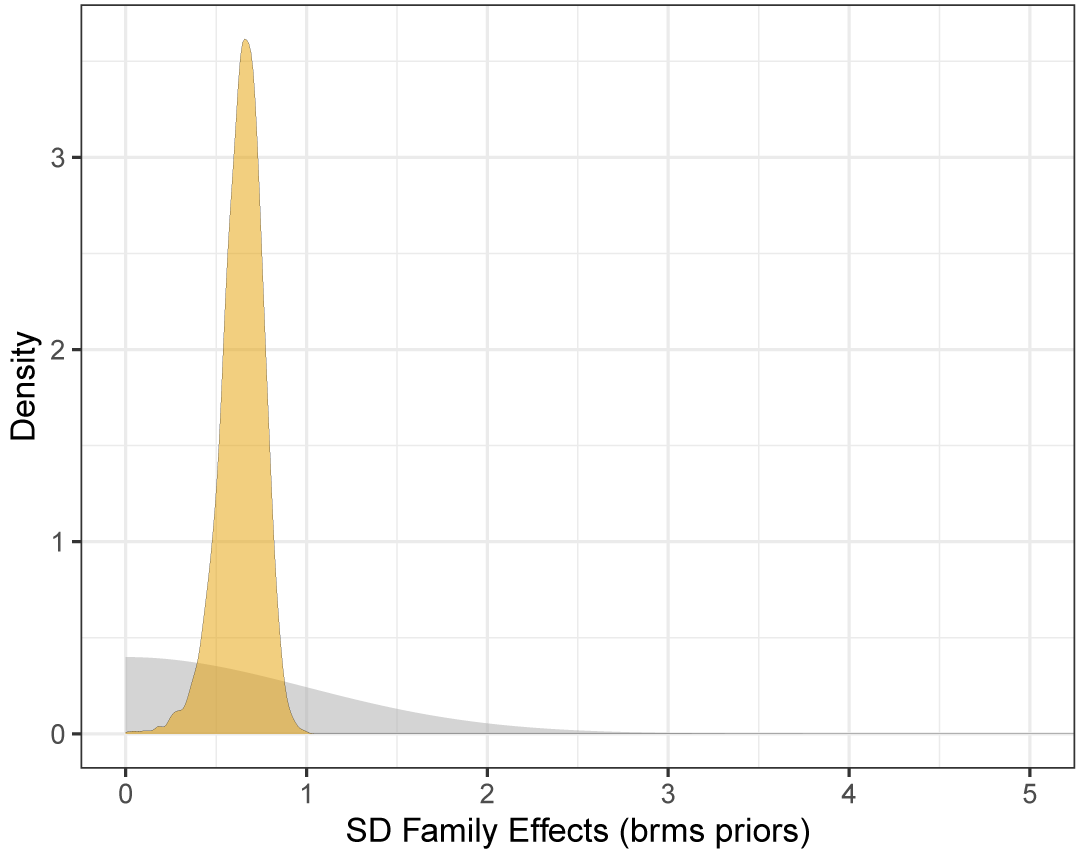
Prior (grey) and posterior (yellow) distributions for the Family Effects standard deviation parameter for the simple rarity model using custom priors.

### 2.3 Full threatened model - IUCN vetted species - *brms* default priors

**Table SM 2:** Summary of model output for *β* and *σ* parameters, including posterior means, posterior standard deviations, 2.5%, 50%, and 97.5% quantiles, the effective sample size (n eff), and the potential scale reduction statistic (Rhat). n=236; RMSE=5.94 (+/-0.72 SD); NRMSE = 0.20 (+/-0.03).

|  | mean | sd | 2.5% | 50% | 97.5% | n_eff | Rhat |
| --- | --- | --- | --- | --- | --- | --- | --- |
| Diversification rate | 0.47 | 0.33 | -0.18 | 0.47 | 1.13 | 4411.32 | 1.00 |
| Famly age | 0.42 | 0.31 | -0.17 | 0.42 | 1.01 | 4175.69 | 1.00 |
| Woodiness | 0.61 | 0.14 | 0.34 | 0.61 | 0.87 | 3874.33 | 1.00 |
| Herbaceous | -1.05 | 0.23 | -1.48 | -1.05 | -0.59 | 2332.31 | 1.00 |
| Climate sum | -0.26 | 0.10 | -0.45 | -0.26 | -0.07 | 4508.21 | 1.00 |
| Number of species hybrids | -0.18 | 0.13 | -0.43 | -0.18 | 0.08 | 4658.36 | 1.00 |
| Hermaphroditic | 0.13 | 0.23 | -0.30 | 0.12 | 0.58 | 4991.91 | 1.00 |
| Dioecious | -0.18 | 0.15 | -0.48 | -0.18 | 0.13 | 4657.14 | 1.00 |
| Monoecious | -0.14 | 0.16 | -0.45 | -0.15 | 0.17 | 4848.15 | 1.00 |
| Asexual | 0.05 | 0.16 | -0.28 | 0.05 | 0.37 | 4352.38 | 1.00 |
| Fleshy fruit | 0.03 | 0.15 | -0.25 | 0.03 | 0.32 | 4339.32 | 1.00 |
| Animal pollinated | 0.55 | 0.25 | 0.05 | 0.55 | 1.04 | 4139.00 | 1.00 |
| SD brownian effects | 0.52 | 0.28 | 0.04 | 0.53 | 1.06 | 472.22 | 1.01 |
| SD family specific effects | 0.73 | 0.12 | 0.47 | 0.74 | 0.92 | 795.47 | 1.01 |

**Figure SM 9:**
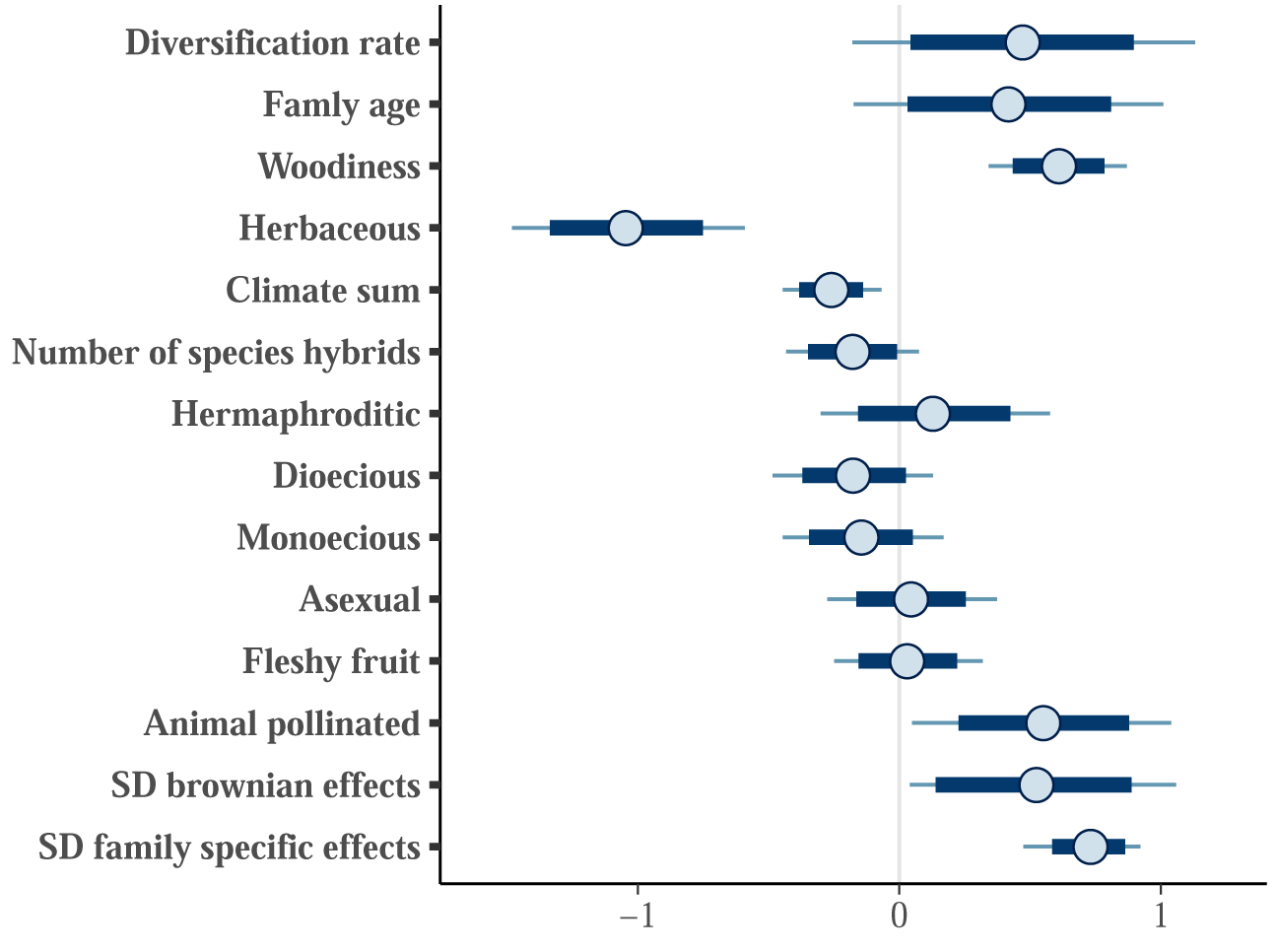
Forest plot for estimated *β* and *σ* parameters for the full threatened model with IUCN vetted species. Points represent posterior means, with thick lines represeting 80% credible intervals, and thin lines representing 95% credible intervals.

**Figure SM 10:**
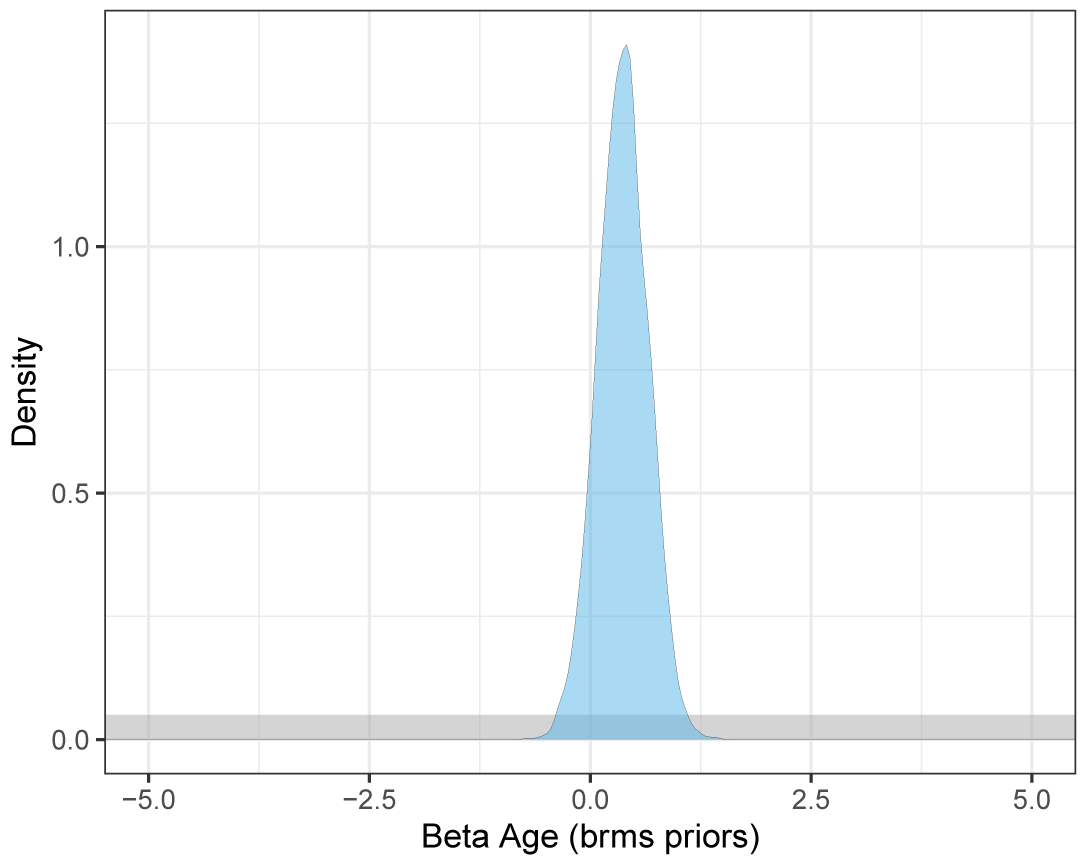
Prior (grey) and posterior (blue) distributions for the Age regression coefficient for the full rarity model using *brms* priors. The improper uniform prior was restricted to (−10,10) to aid visualization.

**Figure SM 11:**
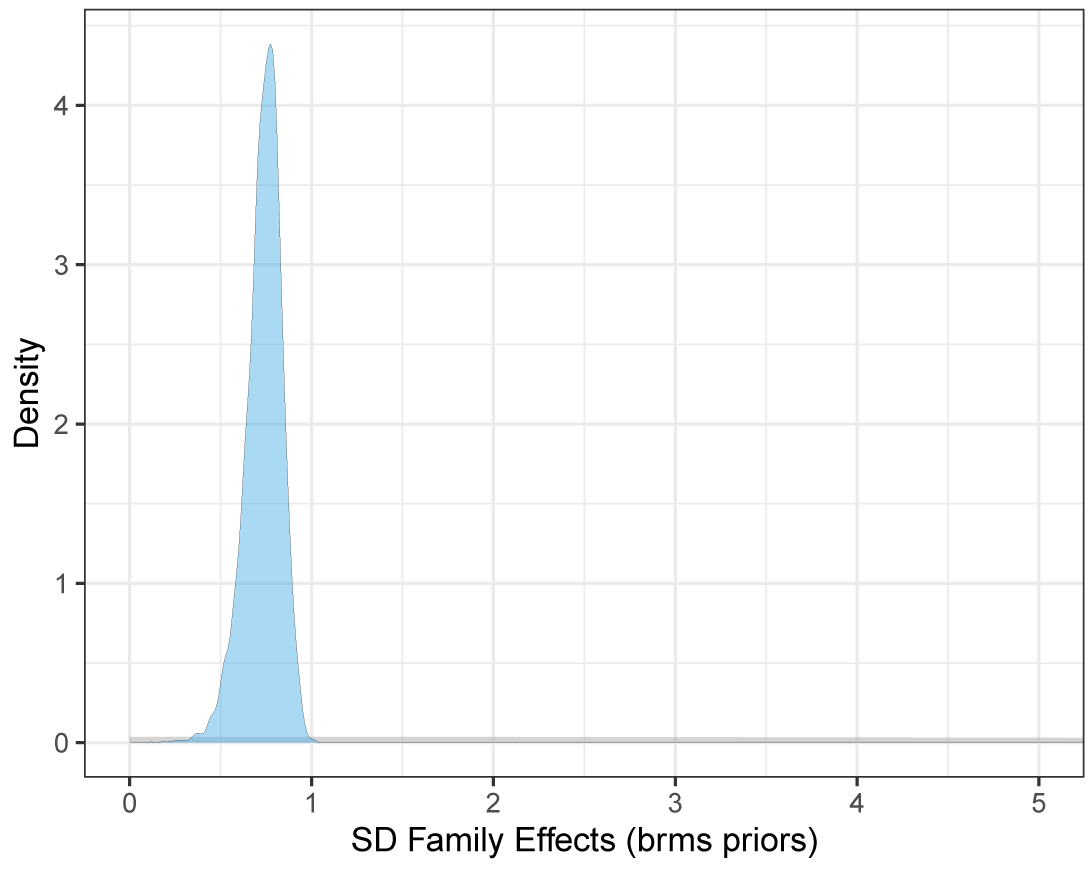
Prior (grey) and posterior (blue) distributions for the Family Effects standard deviation parameter for the full rarity model using *brms* priors.

### 2.4 Full threatened model - IUCN vetted species - custom priors

**Table SM 2:** Summary of model output for *β* and *σ* parameters, including posterior means, posterior standard deviations, 2.5%, 50%, and 97.5% quantiles, the effective sample size (n eff), and the potential scale reduction statistic (Rhat). n=236; RMSE=5.93 (+/-0.73 SD); NRMSE = 0.20 (+/-0.03).

|  | mean | sd | 2.5% | 50% | 97.5% | n_eff | Rhat |
| --- | --- | --- | --- | --- | --- | --- | --- |
| Diversification rate | 0.44 | 0.32 | -0.20 | 0.44 | 1.07 | 4537.10 | 1.00 |
| Famly age | 0.40 | 0.29 | -0.19 | 0.40 | 0.95 | 3913.99 | 1.00 |
| Woodiness | 0.61 | 0.13 | 0.35 | 0.61 | 0.87 | 4766.49 | 1.00 |
| Herbaceous | -1.04 | 0.22 | -1.46 | -1.04 | -0.59 | 3420.69 | 1.00 |
| Climate sum | -0.26 | 0.09 | -0.44 | -0.26 | -0.07 | 4853.71 | 1.00 |
| Number of species hybrids | -0.18 | 0.13 | -0.43 | -0.18 | 0.08 | 5120.11 | 1.00 |
| Hermaphroditic | 0.14 | 0.23 | -0.32 | 0.14 | 0.58 | 4905.70 | 1.00 |
| Dioecious | -0.18 | 0.15 | -0.46 | -0.18 | 0.12 | 4982.72 | 1.00 |
| Monoecious | -0.14 | 0.15 | -0.44 | -0.14 | 0.16 | 4151.91 | 1.00 |
| Asexual | 0.04 | 0.16 | -0.27 | 0.04 | 0.37 | 4815.08 | 1.00 |
| Fleshy fruit | 0.03 | 0.15 | -0.25 | 0.03 | 0.33 | 4621.55 | 1.00 |
| Animal pollinated | 0.54 | 0.25 | 0.04 | 0.54 | 1.02 | 4005.88 | 1.00 |
| SD brownian effects | 0.49 | 0.26 | 0.03 | 0.50 | 1.00 | 798.59 | 1.00 |
| SD family specific effects | 0.74 | 0.11 | 0.50 | 0.75 | 0.92 | 1078.48 | 1.00 |

**Figure SM 12:**
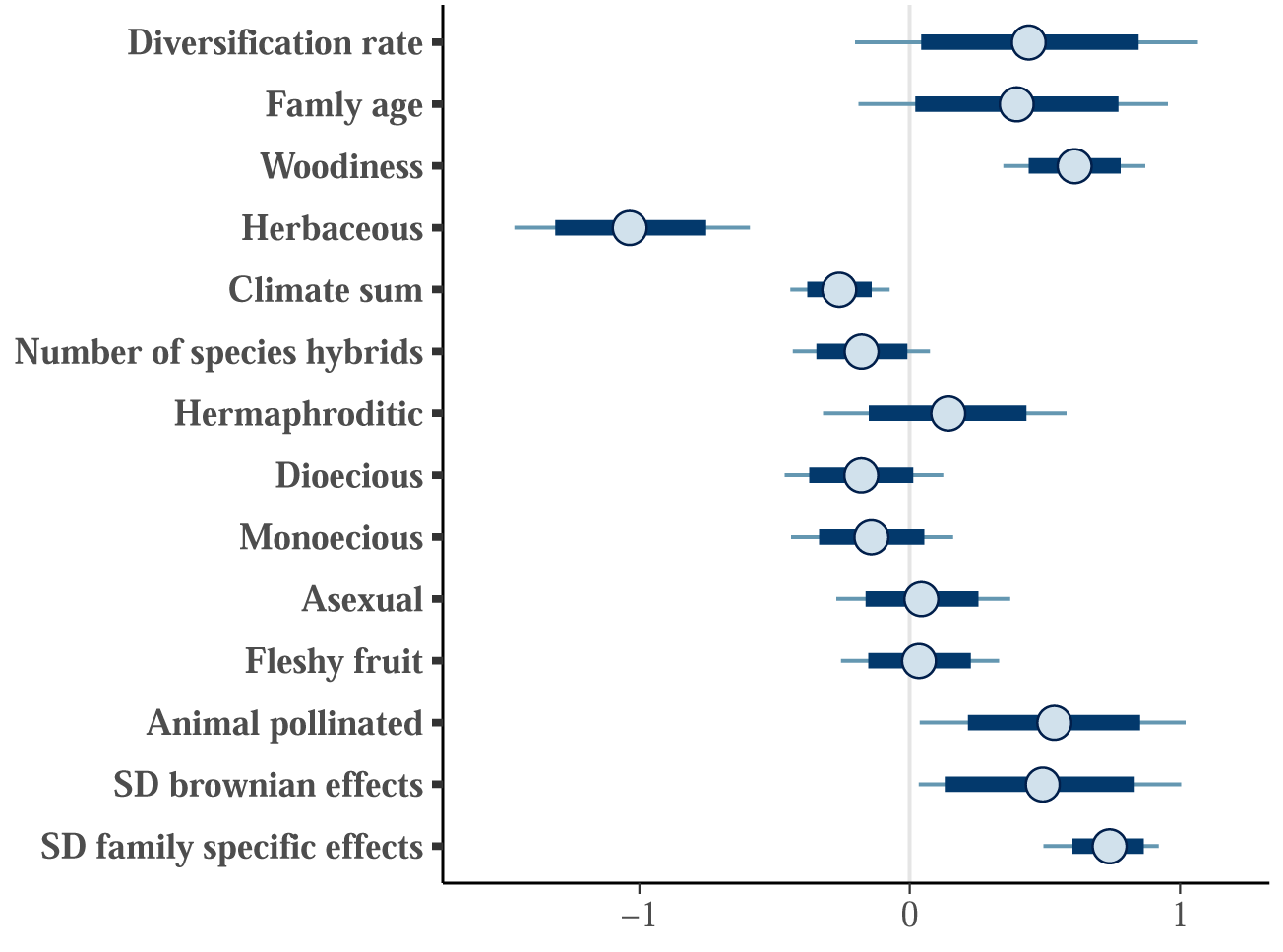
Forest plot for estimated *β* and *σ* parameters for the full threatened model with IUCN vetted species with custom priors. Points represent posterior means, with thick lines represeting 80% credible intervals, and thin lines representing 95% credible intervals.

**Figure SM 13:**
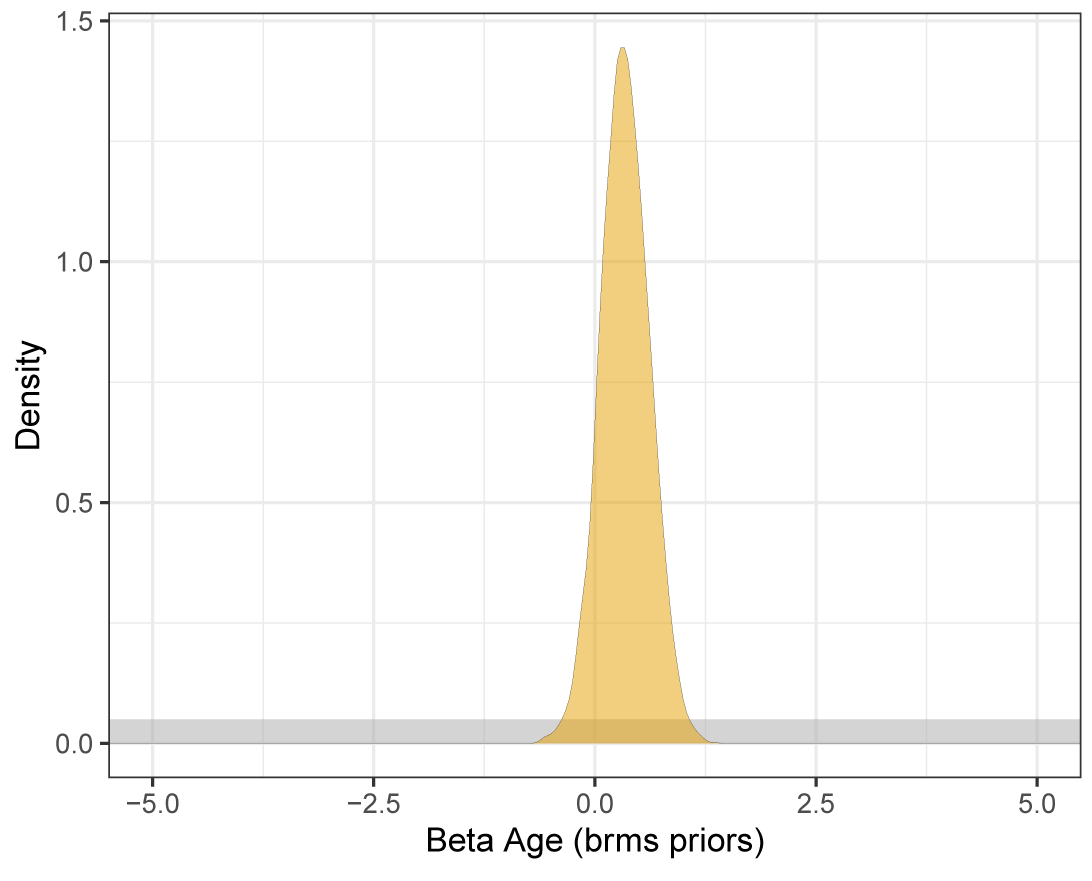
Prior (grey) and posterior (yellow) distributions for the Age regression coefficient for the full rarity model using custom priors. The improper uniform prior was restricted to (−10,10) to aid visualization.

**Figure SM 14:**
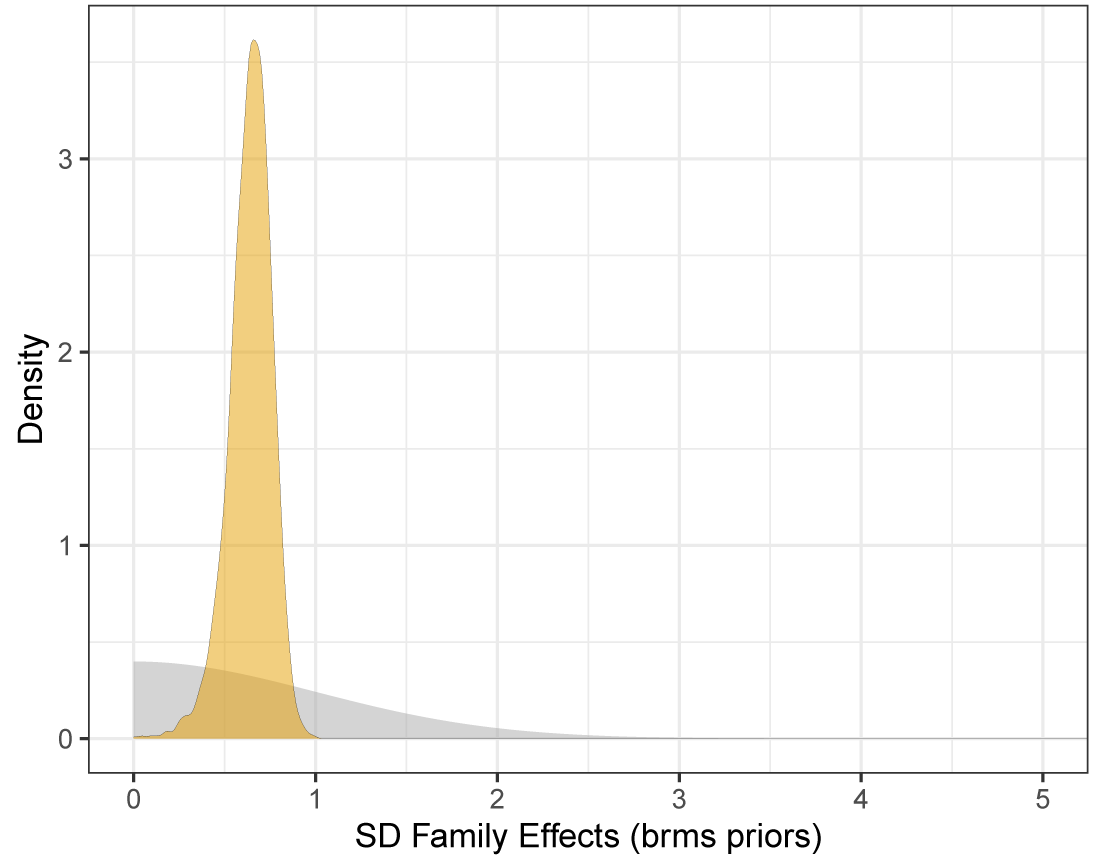
Prior (grey) and posterior (yellow) distributions for the Family Effects standard deviation parameter for the full rarity model using custom priors.

### 2.5 Naturalization model

**Table SM 3:** Summary of model output for *β* and *σ* parameters, including posterior means, posterior standard deviations, 2.5%, 50%, and 97.5% quantiles, the effective sample size (n eff), and the potential scale reduction statistic (Rhat). n=395; RMSE=7.57 (+/-1.08 SD); NRMSE = 0.54 (+/-0.08).

|  | mean | sd | 2.5% | 50% | 97.5% | n_eff | Rhat |
| --- | --- | --- | --- | --- | --- | --- | --- |
| Diversification rate | -1.25 | 0.25 | -1.72 | -1.25 | -0.76 | 4733.68 | 1.00 |
| Famly age | -0.84 | 0.28 | -1.38 | -0.84 | -0.30 | 4829.02 | 1.00 |
| Hybrids | 0.29 | 0.12 | 0.05 | 0.29 | 0.53 | 4517.69 | 1.00 |
| Animal pollinated | -0.56 | 0.26 | -1.06 | -0.56 | -0.05 | 4491.14 | 1.00 |
| Fleshy fruit | -0.01 | 0.14 | -0.28 | -0.00 | 0.26 | 4757.24 | 1.00 |
| Climate sum | 0.33 | 0.09 | 0.14 | 0.33 | 0.51 | 4612.59 | 1.00 |
| Herbaceous | 0.64 | 0.21 | 0.24 | 0.64 | 1.05 | 4822.37 | 1.00 |
| Annual | 0.38 | 0.18 | 0.02 | 0.38 | 0.74 | 4643.58 | 1.00 |
| Axsexual | 0.45 | 0.17 | 0.10 | 0.45 | 0.79 | 4600.48 | 1.00 |
| SD brownian effects | 1.55 | 0.18 | 1.16 | 1.57 | 1.87 | 639.44 | 1.01 |
| SD family specific effects | 0.35 | 0.20 | 0.02 | 0.34 | 0.74 | 419.10 | 1.01 |

**Figure SM 15:**
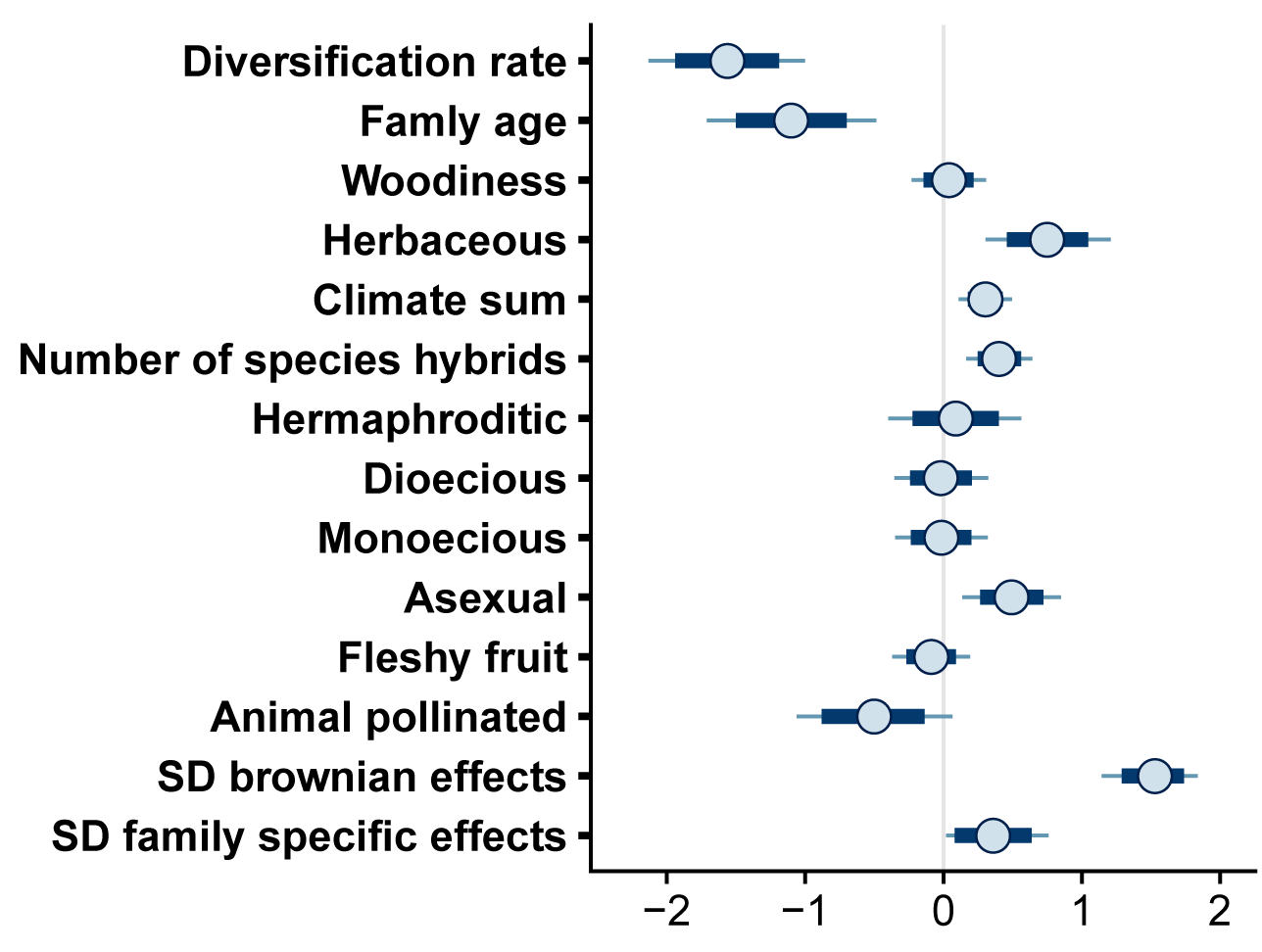
Forest plot for estimated *β* and *σ* parameters for the naturalization model. Points represent posterior means, with thick lines represeting 80% credible intervals, and thin lines representing 95% credible intervals.

### 3 Sensitivity analyses

#### 3.1 Full threatened model: no family effects

**Table SM 4:** Summary of model output for *β* and *σ* parameters, including posterior means, posterior standard deviations, 2.5%, 50%, and 97.5% quantiles, the effective sample size (n eff), and the potential scale reduction statistic (Rhat). n=236; RMSE=16.20 (+/-0.98 SD); NRMSE= 0.55 (+/-0.03).

|  | mean | sd | 2.5% | 50% | 97.5% | n_eff | Rhat |
| --- | --- | --- | --- | --- | --- | --- | --- |
| Diversification rate | 0.18 | 0.09 | -0.00 | 0.18 | 0.36 | 4842.48 | 1.00 |
| Famly age | 0.21 | 0.09 | 0.04 | 0.21 | 0.38 | 4644.13 | 1.00 |
| Woodiness | 0.50 | 0.03 | 0.44 | 0.50 | 0.56 | 5171.53 | 1.00 |
| Herbaceous | -0.71 | 0.05 | -0.81 | -0.71 | -0.61 | 5069.99 | 1.00 |
| Climate sum | -0.37 | 0.02 | -0.41 | -0.37 | -0.32 | 4836.18 | 1.00 |
| Number of species hybrids | -0.15 | 0.04 | -0.24 | -0.15 | -0.07 | 4870.95 | 1.00 |
| Hermaphroditic | 0.19 | 0.06 | 0.07 | 0.19 | 0.31 | 4867.92 | 1.00 |
| Dioecious | 0.12 | 0.03 | 0.05 | 0.12 | 0.19 | 4568.06 | 1.00 |
| Monoecious | -0.26 | 0.04 | -0.34 | -0.26 | -0.18 | 4373.63 | 1.00 |
| Asexual | 0.18 | 0.03 | 0.11 | 0.18 | 0.25 | 4663.13 | 1.00 |
| Fleshy fruit | -0.24 | 0.02 | -0.28 | -0.24 | -0.20 | 4868.31 | 1.00 |
| Animal pollinated | 0.92 | 0.06 | 0.80 | 0.92 | 1.03 | 5010.02 | 1.00 |

**Figure SM 16:**
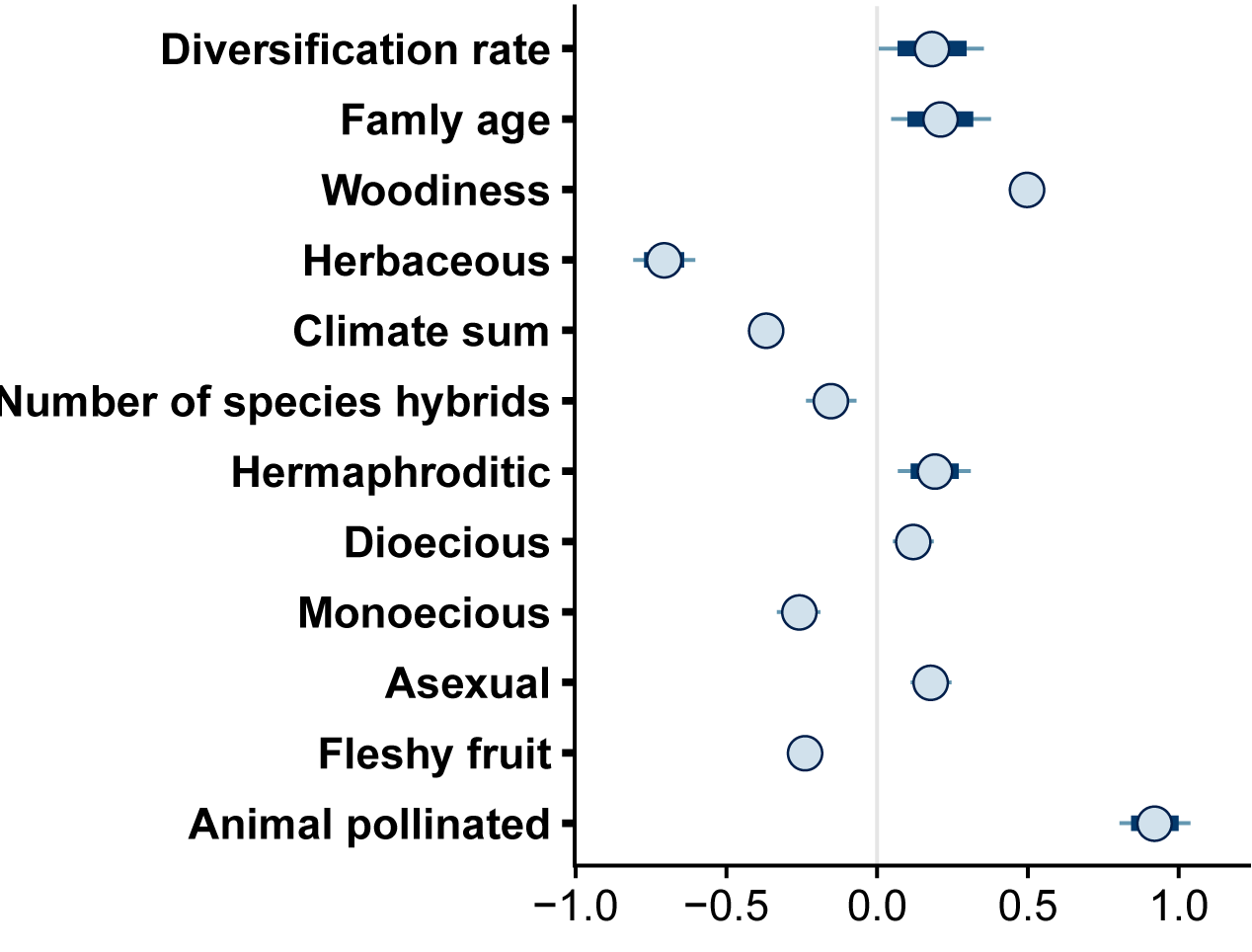
Forest plot for estimated *β* and *σ* parameters for the threatened model with no family effects. Points represent posterior means, with thick lines represeting 80% credible intervals, and thin lines representing 95% credible intervals.

### 3.2 Full threatened model: only non-Brownian family effects

**Table SM 5:** Summary of model output for *β* and *σ* parameters, including posterior means, posterior standard deviations, 2.5%, 50%, and 97.5% quantiles, the effective sample size (n eff), and the potential scale reduction statistic (Rhat). n=236; RMSE=5.94 (+/-0.71 SD); NRMSE = 0.20 (+/-0.02).

|  | mean | sd | 2.5% | 50% | 97.5% | n_eff | Rhat |
| --- | --- | --- | --- | --- | --- | --- | --- |
| Diversification rate | 0.47 | 0.32 | -0.17 | 0.47 | 1.11 | 3299.25 | 1.00 |
| Family age | 0.49 | 0.29 | -0.06 | 0.49 | 1.05 | 4013.44 | 1.00 |
| Woodiness | 0.65 | 0.12 | 0.41 | 0.65 | 0.88 | 3314.83 | 1.00 |
| Herbaceous | -1.17 | 0.19 | -1.55 | -1.17 | -0.80 | 3679.59 | 1.00 |
| Climate sum | -0.27 | 0.09 | -0.45 | -0.27 | -0.08 | 3554.38 | 1.00 |
| Number of species hybrids | -0.17 | 0.13 | -0.43 | -0.17 | 0.09 | 4490.73 | 1.00 |
| Hermaphroditic | 0.13 | 0.23 | -0.33 | 0.13 | 0.56 | 3460.29 | 1.00 |
| Dioecious | -0.18 | 0.16 | -0.47 | -0.18 | 0.13 | 3374.37 | 1.00 |
| Monoecious | -0.15 | 0.16 | -0.46 | -0.15 | 0.16 | 3365.90 | 1.00 |
| Asexual | 0.06 | 0.16 | -0.26 | 0.06 | 0.37 | 3226.05 | 1.00 |
| Fleshy fruit | 0.04 | 0.15 | -0.25 | 0.05 | 0.34 | 4319.92 | 1.00 |
| Animal pollinated | 0.63 | 0.24 | 0.16 | 0.63 | 1.10 | 4088.85 | 1.00 |
| SD family specific effects | 0.84 | 0.06 | 0.73 | 0.84 | 0.97 | 3310.40 | 1.00 |

**Figure SM 17:**
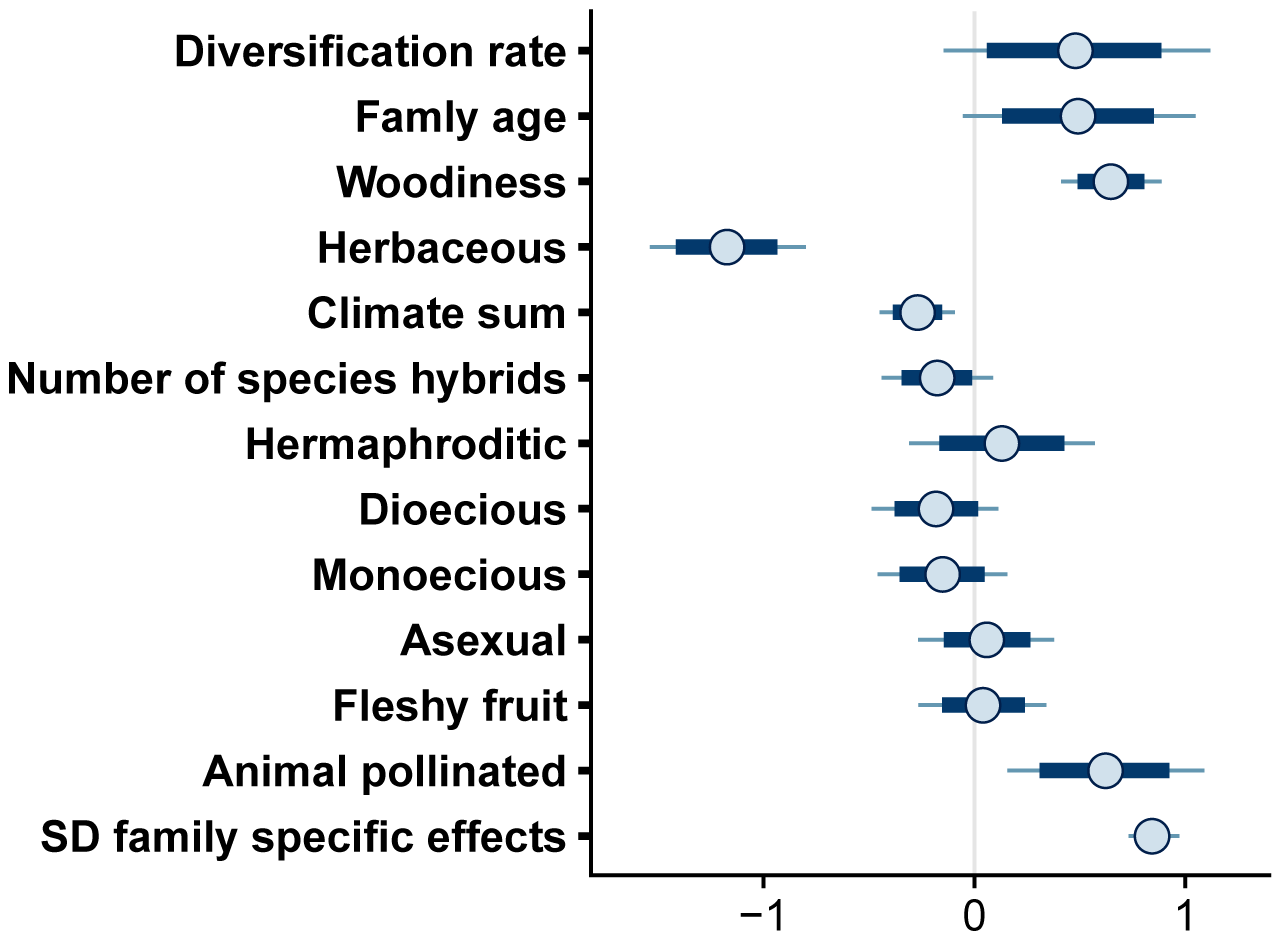
Forest plot for estimated *β* and *σ* parameters for the threatened model with non-Brownian family effects. Points represent posterior means, with thick lines represeting 80% credible intervals, and thin lines representing 95% credible intervals.

### 3.3 Full threatened model: only Brownian family effects

**Table SM 6:** Summary of model output for *β* and *σ* parameters, including posterior means, posterior standard deviations, 2.5%, 50%, and 97.5% quantiles, the effective sample size (n eff), and the potential scale reduction statistic (Rhat). n=236; RMSE=5.94 (+/-0.71 SD); NRMSE = 0.20 (+/-0.02).

|  | mean | sd | 2.5% | 50% | 97.5% | n_eff | Rhat |
| --- | --- | --- | --- | --- | --- | --- | --- |
| Diversification rate | 0.55 | 0.33 | -0.12 | 0.55 | 1.18 | 3231.76 | 1.00 |
| Famly age | 0.35 | 0.33 | -0.31 | 0.35 | 0.99 | 2949.88 | 1.00 |
| Woodiness | 0.57 | 0.14 | 0.30 | 0.57 | 0.84 | 2759.86 | 1.00 |
| Herbaceous | -0.89 | 0.23 | -1.35 | -0.89 | -0.44 | 2846.98 | 1.00 |
| Climate sum | -0.24 | 0.09 | -0.42 | -0.24 | -0.05 | 3148.80 | 1.00 |
| Number of species hybrids | -0.19 | 0.13 | -0.44 | -0.18 | 0.06 | 4371.60 | 1.00 |
| Hermaphroditic | 0.10 | 0.22 | -0.34 | 0.10 | 0.54 | 3358.10 | 1.00 |
| Dioecious | -0.19 | 0.15 | -0.49 | -0.19 | 0.09 | 3263.29 | 1.00 |
| Monoecious | -0.09 | 0.15 | -0.38 | -0.10 | 0.20 | 3056.87 | 1.00 |
| Asexual | 0.01 | 0.16 | -0.30 | 0.01 | 0.32 | 2792.53 | 1.00 |
| Fleshy fruit | -0.08 | 0.12 | -0.32 | -0.08 | 0.17 | 4134.48 | 1.00 |
| Animal pollinated | 0.44 | 0.25 | -0.04 | 0.44 | 0.94 | 3792.81 | 1.00 |
| SD brownian effects | 1.31 | 0.10 | 1.13 | 1.31 | 1.51 | 3628.27 | 1.00 |

**Figure SM 18:**
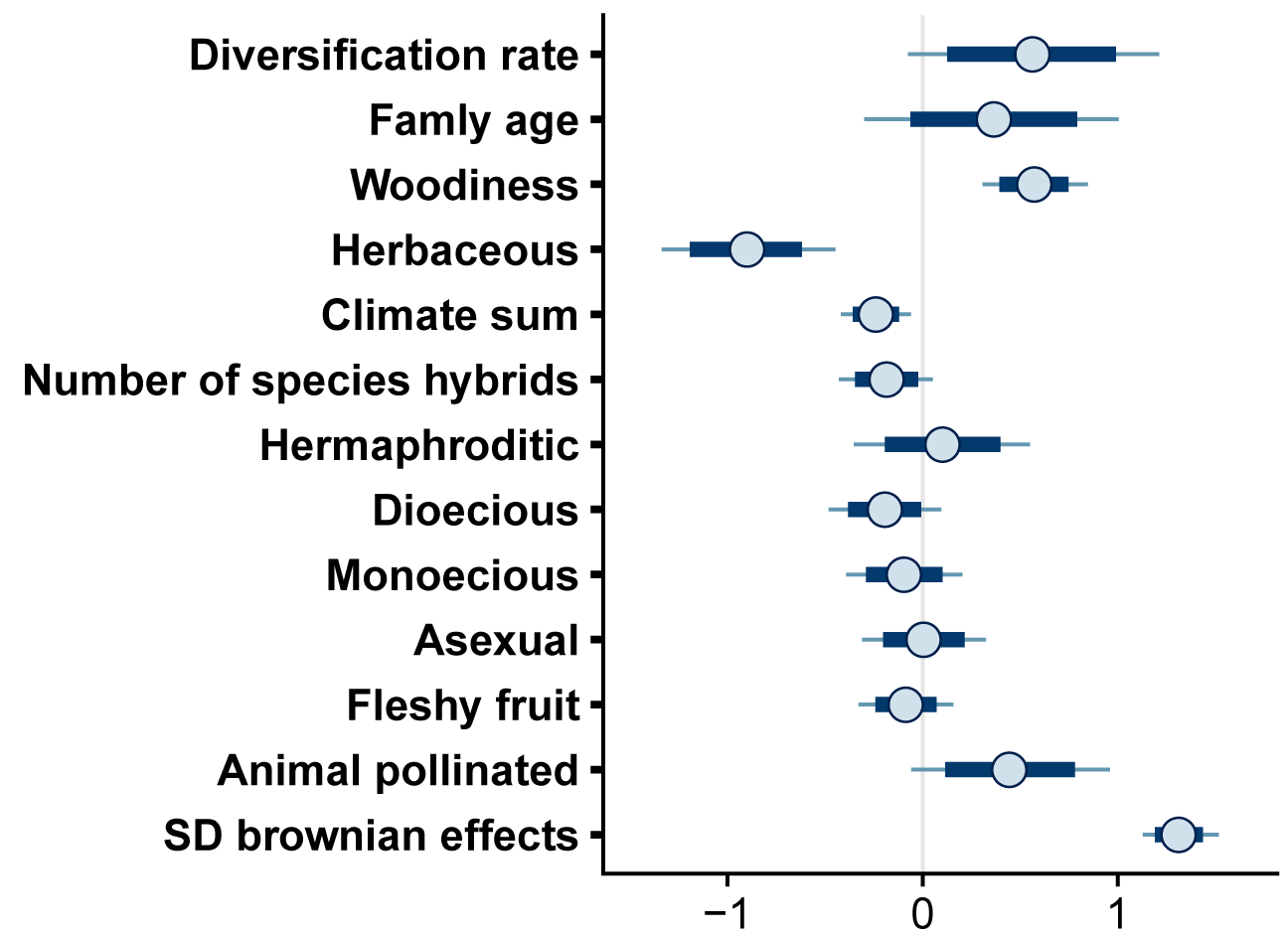
Forest plot for estimated *β* and *σ* parameters for the threatened model with Brownian family effects. Points represent posterior means, with thick lines represeting 80% credible intervals, and thin lines representing 95% credible intervals.

### 3.4 Full threatened model: including non-IUCN vetted species Invasive model

**Table SM 7:** Summary of model output for *β* and *σ* parameters, including posterior means, posterior standard deviations, 2.5%, 50%, and 97.5% quantiles, the effective sample size (n eff), and the potential scale reduction statistic (Rhat). n=395; RMSE=6.82 (+/-0.76 SD); NRMSE = 0.49 (+/-0.05).

3.4 Full threatened model: including non-IUCN vetted species
|  | mean | sd | 2.5% | 50% | 97.5% | n_eff | Rhat |
| --- | --- | --- | --- | --- | --- | --- | --- |
| Diversification rate | 0.69 | 0.36 | 0.00 | 0.69 | 1.40 | 4087.61 | 1.00 |
| Famly age | 0.51 | 0.32 | -0.10 | 0.51 | 1.15 | 4584.84 | 1.00 |
| Woodiness | -0.41 | 0.16 | -0.73 | -0.41 | -0.10 | 3899.62 | 1.00 |
| Herbaceous | -0.62 | 0.26 | -1.12 | -0.62 | -0.13 | 2609.26 | 1.00 |
| Climate sum | -0.18 | 0.11 | -0.40 | -0.18 | 0.03 | 4592.67 | 1.00 |
| Number of species hybrids | 0.03 | 0.16 | -0.28 | 0.02 | 0.34 | 3696.19 | 1.00 |
| Hermaphroditic | -0.11 | 0.27 | -0.64 | -0.12 | 0.41 | 4531.19 | 1.00 |
| Dioecious | -0.19 | 0.19 | -0.55 | -0.19 | 0.16 | 4460.67 | 1.00 |
| Monoecious | 0.07 | 0.18 | -0.29 | 0.07 | 0.44 | 4861.42 | 1.00 |
| Asexual | 0.15 | 0.21 | -0.25 | 0.16 | 0.56 | 4629.34 | 1.00 |
| Fleshy fruit | 0.24 | 0.18 | -0.12 | 0.24 | 0.60 | 5103.72 | 1.00 |
| Animal pollinated | 0.17 | 0.31 | -0.41 | 0.17 | 0.78 | 4352.88 | 1.00 |
| SD brownian effects | 0.68 | 0.32 | 0.07 | 0.68 | 1.33 | 532.46 | 1.01 |
| SD family specific effects | 1.07 | 0.13 | 0.78 | 1.08 | 1.29 | 752.64 | 1.00 |

**Figure SM 19:**
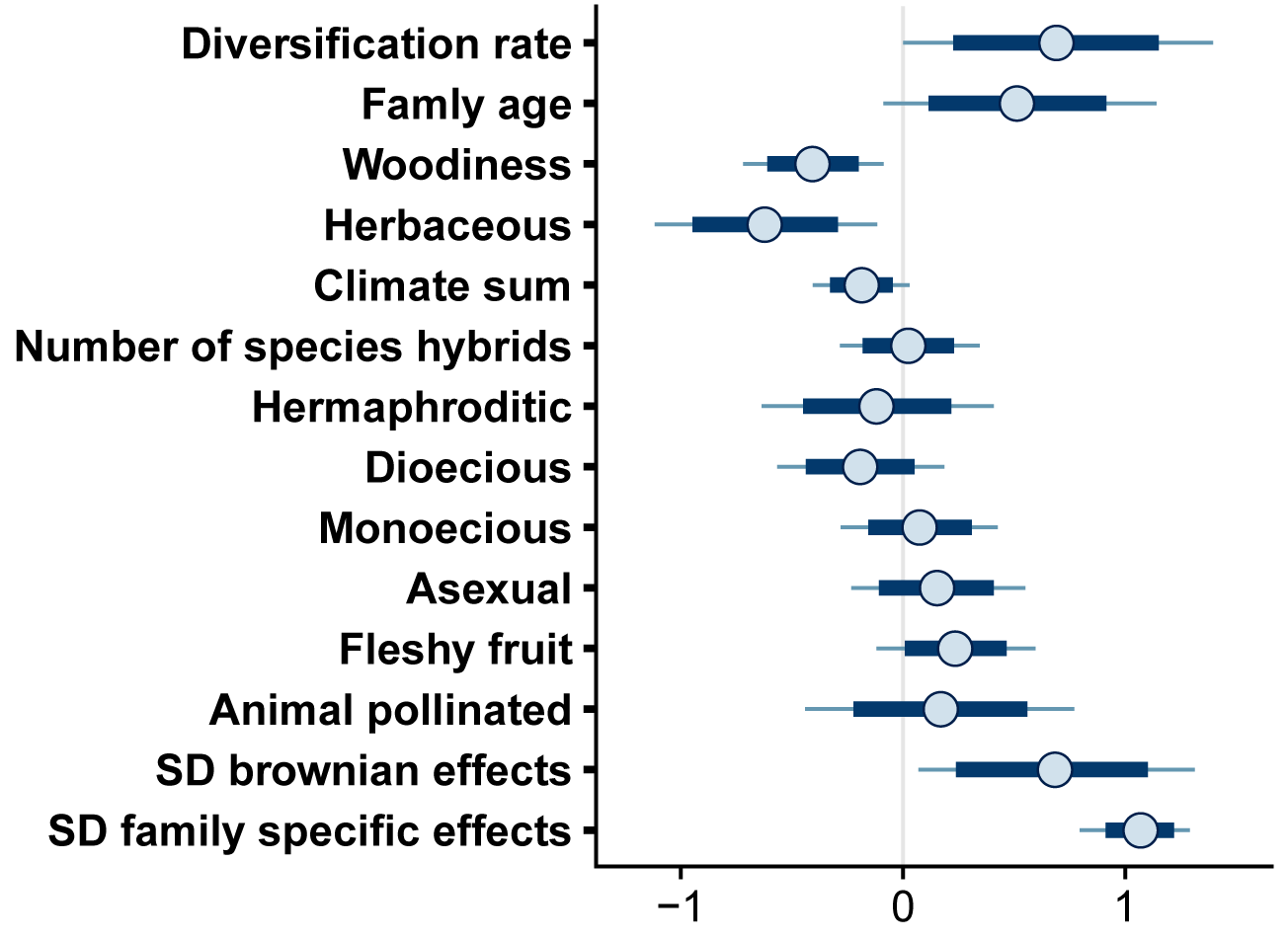
Forest plot for estimated *β* and *σ* parameters for the threatened model with non-IUCN vetted species. Points represent posterior means, with thick lines represeting 80% credible intervals, and thin lines representing 95% credible intervals.

### Invasive model

**Table SM 8:** Summary of model output for *β* and *σ* parameters, including posterior means, posterior standard deviations, 2.5%, 50%, and 97.5% quantiles, the effective sample size (n eff), and the potential scale reduction statistic (Rhat). n=395; RMSE=4.68 (+/-0.66 SD); NRMSE = 0.78 (+/-0.11).

3.5 Invasive model
|  | mean | sd | 2.5% | 50% | 97.5% | n_eff | Rhat |
| --- | --- | --- | --- | --- | --- | --- | --- |
| Diversification rate | -0.97 | 0.31 | -1.56 | -0.98 | -0.33 | 4530.29 | 1.00 |
| Famly age | -0.76 | 0.35 | -1.46 | -0.76 | -0.09 | 4119.70 | 1.00 |
| Hybrids | 0.35 | 0.15 | 0.05 | 0.35 | 0.63 | 4144.23 | 1.00 |
| Animal pollinated | -0.42 | 0.32 | -1.04 | -0.42 | 0.21 | 3702.58 | 1.00 |
| Fleshy fruit | 0.07 | 0.16 | -0.25 | 0.07 | 0.39 | 4626.87 | 1.00 |
| Climate sum | 0.18 | 0.11 | -0.04 | 0.18 | 0.41 | 4319.14 | 1.00 |
| Herbaceous | 0.66 | 0.25 | 0.18 | 0.66 | 1.15 | 4057.81 | 1.00 |
| Annual | 0.22 | 0.21 | -0.19 | 0.21 | 0.63 | 4291.63 | 1.00 |
| Axsexual | 0.42 | 0.21 | 0.01 | 0.43 | 0.82 | 3729.91 | 1.00 |
| SD brownian effects | 1.69 | 0.22 | 1.20 | 1.71 | 2.07 | 677.12 | 1.01 |
| SD family specific effects | 0.40 | 0.25 | 0.02 | 0.40 | 0.89 | 400.47 | 1.01 |

**Figure SM 20:**
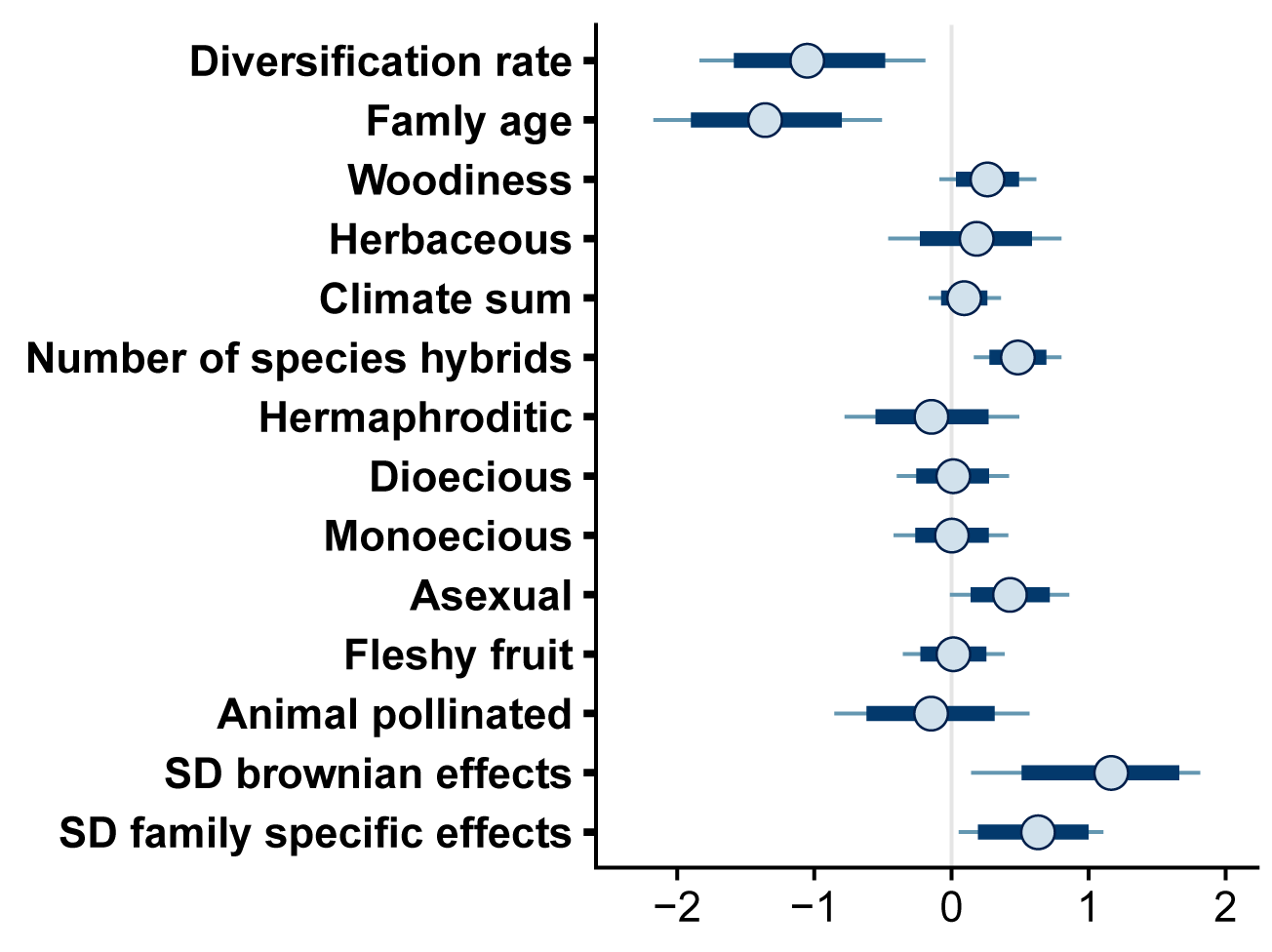
Forest plot for estimated *β* and *σ* parameters for the invasive model. Points represent posterior means, with thick lines represeting 80% credible intervals, and thin lines representing 95% credible intervals.

## Notes

### Competing Interest Statement

The authors have declared no competing interest.

